# Which plant traits increase soil carbon sequestration? Empirical evidence from a long-term poplar genetic diversity trial

**DOI:** 10.1101/2025.02.17.638464

**Authors:** John L. Field, Brandon P. Sloan, Matthew E. Craig, Parker Calloway, Sarah L. Ottinger, Thomas Mead, Rose Z. Abramoff, Mirko Pavicic Venegas, Hari B. Chhetri, Kathy Haiby, Udaya C. Kalluri, Wellington Muchero, Christopher W. Schadt, Melanie A. Mayes

**Author notes:** Notice: This manuscript has been authored by UT-Battelle, LLC, under contract DE-AC05-00OR22725 with the US Department of Energy (DOE). The US government retains and the publisher, by accepting the work for publication, acknowledges that the US government retains a non-exclusive, paid-up, irrevocable, world-wide license to publish or reproduce the submitted manuscript version of this work, or allow others to do so, for US government purposes. DOE will provide public access to these results of federally sponsored research in accordance with the DOE Public Access Plan (https://energy.gov/doe-public-access-plan).

## Abstract

Breeding or engineering crops to increase soil organic carbon (SOC) storage is a potential route to land-based carbon dioxide removal on working agricultural lands. However, due to limited observational datasets plus shifting paradigms of SOC stabilization, there is a lack of understanding of which plant traits to target for SOC enhancement or the ultimate sequestration potential of such measures. Existing long-term common gardens of genetically diverse plant populations may provide an opportunity to evaluate biological controls on SOC outcomes, separate from environmental or management variability. Here we report on soil carbon and root chemical data collected for 24 genotypes within a 13-year-old common garden in northwestern Oregon planted with over a thousand natural variants of Populus trichocarpa. Fractionating surface soil (0–15 cm) revealed substantial variation in stocks of mineral-associated organic matter (MAOM; 18–67 tonnes C/ha) and particulate organic matter (POM; 2–22 tonnes C/ha). Tree genotype explained 24 and 26% of the MAOM and POM stock variability, respectively, after controlling for background variability. We found minimal association between SOC concentration and either aboveground tree productivity or conventional metrics of root biomass recalcitrance (C/N ratios and lignin content). However, root elemental content appeared influential for MAOM-C, which showed a strong positive association with root aluminum (Al), and strong negative association with root boron (B) and magnesium (Mg). Furthermore, root concentrations of these elements were highly heritable (57–78%) and not simply a reflection of background variation in bulk soil elemental concentrations. We estimate that surface SOC stocks under these 24 genotypes have diverged at rates of up to 1.2–4.3 tonnes C/ha/yr. These results suggest that genetic diversity trials have value for elucidating biological controls on SOC dynamics, and that traits associated with root elemental content may be an important target for enhancing SOC.

## 1 Introduction

Global decarbonization scenarios project a need for 5–180 gigatons of carbon removal (Gt C) from the atmosphere by the end of the century (IPCC, 2022). Historically, human agricultural practices are responsible for 116 Gt C that has been lost from soils to the atmosphere (Sanderman et al., 2017), compared to 500 Gt C of cumulative fossil fuel emissions to date (Friedlingstein et al., 2024). Reversing these losses and restoring agricultural soils towards or beyond their natural soil organic carbon (SOC) storage levels is a carbon removal opportunity with substantial agricultural and environmental co-benefits. The universal adoption of agricultural conservation management best practices such as tillage intensity reduction, cover-cropping, and conversion of degraded agricultural land to perennial crops has the potential to remove 1–3 Gt C/yr from the atmosphere globally (National Academies of Sciences, Engineering, and Medicine, 2019; Paustian, Larson, et al., 2019), while other “frontier” methods (i.e., methods that have not yet been applied at scale due to technical or economic barriers) could plausibly double this removal potential (Paustian, Larson, et al., 2019).

One such frontier method is to breed or engineer agricultural crops (e.g., annual food crops, pasture grasses, or biomass crops) for traits that would lead to enhanced “biosequestration” of SOC (U.S. DOE, 2008) beyond what is possible with current commercial crop varieties and best management practices (National Academies of Sciences, Engineering, and Medicine, 2019; Paustian, Larson, et al., 2019), an approach sometimes termed “carbon farming” (Jansson et al., 2021). There is particular research interest in optimizing biosequestration potential under dedicated perennial energy crops (Jansson et al., 2010), which are projected to play a large role in future decarbonization plans (Butnar et al., 2020). Various plant trait targets have been proposed for enhancing SOC biosequestration (Bardgett et al., 2014; Fierer & Walsh, 2023; Jansson et al., 2010, 2021; Kalluri et al., 2020; Poirier et al., 2018; Wang & Kuzyakov, 2023; Yang et al., 2021). However, there are few studies quantifying the potential impact of these traits in different agricultural systems and environments beyond back of the envelope calculations and some exploratory modeling work (e.g., Cotrufo et al., 2024; Paustian, Campbell, et al., 2016).

There have recently been several major paradigm shifts around soil organic matter (SOM) dynamics and the plant traits most likely to influence them. First, while much past research has focused on SOM formation from aboveground litter, newer work suggests that roots and other belowground inputs are stabilized into SOC with much higher efficiency and thus play an outsized role compared to aboveground inputs (Berhongaray et al., 2019; Sokol & Bradford, 2019). Such belowground inputs are particularly important for bioenergy crops, since aboveground biomass is periodically harvested and exported from the system. Second, while SOM was previously thought to be composed of chemically-recalcitrant plant tissues and microbial by-products inherently resistant to further decomposition, it is now known that stable organic matter in mineral soils mostly consists of simple compounds that have been spared from further microbial processing due to physical separation, chemical protection, or functional complexity in microbe–substrate interactions (Lehmann et al., 2020; Lehmann & Kleber, 2015; Schmidt et al., 2011). Similarly-new paradigms for the classification of different organic matter pools have also increased in importance. Particulate organic matter (POM) is made up of partially-decomposed plant tissues that can be physically protected from soil microbes in microaggregates or soil pore structures (Lavallee et al., 2020), whereas mineral-associated organic matter (MAOM) consists of microbial necromass or simple plant-derived compounds adsorbed to clay minerals (Cotrufo et al., 2022; Rowley et al., 2018; Sokol et al., 2019) or adsorbed to and co-precipitated with iron, manganese, and aluminum (oxyhydr)oxides (Rasmussen et al., 2018; Zhao et al., 2016, 2020). Finally, in addition to being a biomass source for SOM formation, plants are also known affect the decay of existing SOM via priming (Poirier et al., 2018). In particular, low molecular weight compounds exuded by plant roots can exchange with clay surface adsorbed compounds to destabilize existing MAOM, or indirectly stimulate greater microbial decomposition of existing organic matter stocks (Jilling et al., 2021; Keiluweit et al., 2015).

Under the older paradigms, strategies for enhancing biosequestration included selection of plants with more chemically-recalcitrant biomass, i.e., higher C/N ratios, lignin, suberin, or phytoliths (Harman-Ware, Sparks, et al., 2021; Parton et al., 1987; Poirier et al., 2018; Yang et al., 2021). In contrast, the newer paradigms imply that the opposite strategy may be more effective—i.e., that higher-quality biomass can be processed more efficiently by microbes and result in greater levels of microbial biomass, which in turn then leads to more durable MAOM storage (Cotrufo et al., 2013; Poirier et al., 2018). Such approaches have been confirmed via both short-term soil incubations (Ridgeway et al., 2022) and plant microcosm studies (Rossi et al., 2020). MAOM formation might also be increased through selecting plants with high rates of root exudation and rhizodeposition (Jansson et al., 2021; Poirier et al., 2018; Wang & Kuzyakov, 2023), whereas POM might potentially be increased via fast fine root turnover, recalcitrant biomass chemistry, or high levels of root mucilage, exudation, or mycorrhizal association that promote increased soil aggregation (Beattie et al., 2024; Fierer & Walsh, 2023; Jansson et al., 2021; Lynd et al., 2024; Poirier et al., 2018; Wang & Kuzyakov, 2023).

Understanding of SOM drivers has been advanced using data from agricultural or ecological monitoring networks, soil survey databases, and meta-analyses (Hansen et al., 2024; Heckman et al., 2023; King et al., 2023; Lugato et al., 2021) that are rich in environmental variability, but often contain only limited representation of plant functional traits beyond primary production. In contrast, long-term (decade or older) common garden experiments—i.e., multiple species or varieties grown together at the same site—offer the potential to disentangle the relative importance of myriad plant traits on soil carbon levels without being swamped by environmental variability or confounding between plant traits and environmental conditions (Poirier et al., 2018; Waring et al., 2022). Such studies provide mixed evidence of inter-species or inter-variety effects on SOC levels. Long-term common gardens with different tree species grown in monoclonal stands have demonstrated differences in litter layer carbon stocks (Vesterdal et al., 2008) in one experiment, and differences in soil pH and cation content (Reich et al., 2005) and SOM decomposition potential (Hobbie et al., 2007) but not carbon stocks in the mineral soil horizons (Mueller et al., 2012) in another experiment. Pregitzer et al. (2013) sampled individual trees from among 5 genotypes of *Populus angustifolia* planted in three long-term polyclonal grid-planted common gardens in Utah, finding a correlation between tree size and SOC, but no consistent genotype effect. Waring et al. (2022) sampled a 9-year-old common garden with monoclonal plots of 19 different tree species, finding significant differences in soil bulk density and carbon stocks, the latter positively associated with tree productivity but negatively associated with litter recalcitrance. Similar long-term common garden studies of perennial bioenergy grasses have shown species- and cultivar-level differences in root morphology and clay-associated SOC (Kelly-Slatten et al., 2023), though such cultivar-level SOC differences are generally not observed in shorter-duration studies (Adkins et al., 2019; Collins et al., 2010).

In this work, we sampled soil and root biomass from individual trees of 24 different genotypes established in a long-term (13-year-old) polyclonal grid-planted poplar common garden in Oregon, fractionated the 0–15 cm depth soil samples into MAOM and POM, and performed statistical modeling to quantify broad-sense heritability and phenotype effects on soil C levels. Our goals were to a) assess whether substantial differences in SOC are apparent across this site, b) explore associations between plant traits and SOC differences with respect to competing stabilization paradigms, and c) quantify whether those trait and SOC differences are influenced by tree genotype. This work builds on a previous exploratory study that found evidence of genotype-driven differences in soil pH, base cations, and bulk density across this common garden (Craig et al., 2024), and it is meant to support the identification of high-priority trait targets and eventual quantification of the upper potential of enhanced SOC biosequestration for a promising, second-generation bioenergy feedstock.

## 2 Methods

### 2.1 GWAS Population and Experimental Site

The Center for Bioenergy Innovation, GreenWood Resources Inc., and the University of British Columbia have assembled a diverse research population of *Populus trichocarpa* (commonly known as black cottonwood) to support genome-wide association studies (GWAS). This GWAS population includes more than 1,100 unique genotypes collected across the western U.S. and British Columbia (Fig. S1). *P. trichocarpa* is diecious species (thus an obligate outbreeder) that has not previously been domesticated, so this species and research population feature high genetic and trait diversity. This GWAS population been replicated in a number of common garden plantings and studies to explore the genetic controls of biomass yield and phenology (Evans et al., 2014), tissue lignin content (Xie et al., 2018), the associated soil microbiome (Schadt et al., 2024; Veach et al., 2019), and other favorable properties for bioenergy production (Happs et al., 2021).

One of these GWAS common garden sites was established in April 2009 near Clatskanie, Oregon (46°07’16”N, 123°16’13”W; Fig. 1A). The site is on diked former floodplains of the Columbia River, and experiences warm wet summers and cool dry winters. It is mapped in the Soil Survey Geographic (SSURGO) database (Ernstrom & Lytle, 1993) as silt loam from the Wauna and Locoda series, though our own texture measurements show silty clay and silty clay loam in the 0–15 cm surface layer (Fig. S2). The soils are rated as deep, poorly drained, strongly acidic, and exhibiting redox mottling suggestive of significant fluctuation between anaerobic and aerobic conditions (National Cooperative Soil Survey, USDA, 2005).

**Figure 1.**
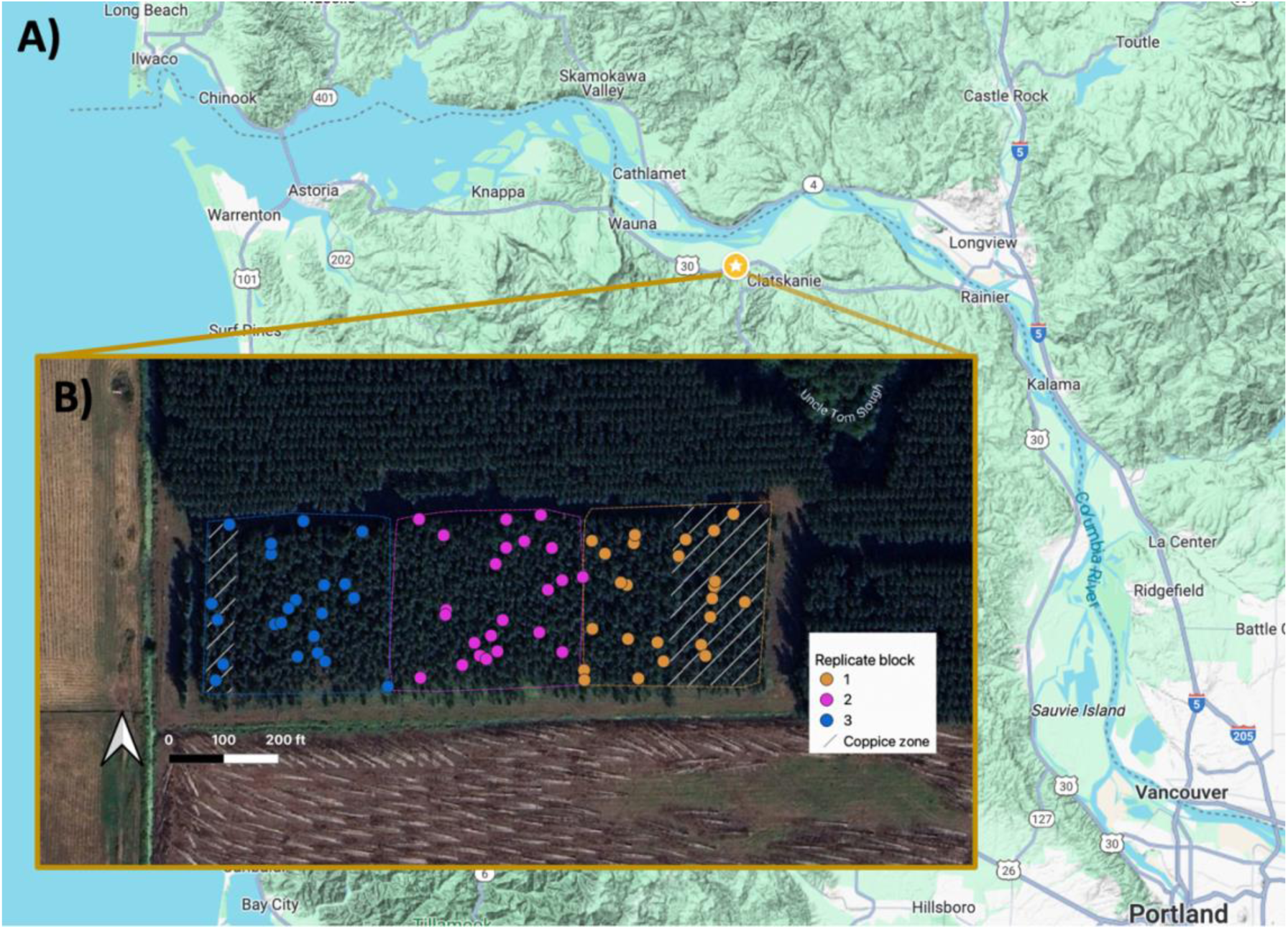
A) Location of poplar common garden site near Clatskanie, in northwest Oregon. B) Inset map showing the 69 individual trees sampled in this work, representing 24 different genotypes replicated across three spatial blocks. Grey hatching at eastern and western parts of blocks 1 and 3 denotes areas that were coppiced in 2012.

Prior to the GWAS planting, commercial hybrid poplar (*P*. *trichocarpa* × *P*. *deltoides*) was grown on the site for approximately 20 years, with the most recent rotation from 1995–2007. The different varieties of *P. trichocarpa* were established in a polyclonal gridded planting, with each genotype replicated across three spatial blocks, and trees completely randomized within each block (Fig. 1B). The planting grid has 3 m by 3 m row spacing (*c.* 1,100 trees per hectare) for a total plot area of 3.1 hectares, with an additional two-tree buffer rows surrounding the plot. In 2012, the aboveground portion of *∼* 1/2 of the easternmost experimental Block 1 and *∼* 1/5 of the westernmost Block 3 were coppiced (Fig. 1B) and allowed to regrow, thus all the trees were functionally either 10 or 13 years old at the time of this study. At the time of our initial study design, 449 of the genotypes had all three of their spatial replicate trees still alive.

### 2.2 Soil and Root Sampling

Of the >1,100 *P. trichocarpa* genotypes originally established in this common garden, we down- selected 24 genotypes for soil and root sampling based on survival of all three tree replicates, and extreme values of foliar chemistry that might affect litter recalcitrance or soil microbial activity. Specifically, we selected for extreme values of ferulic and para-coumaric acid (Tuskan et al., 2019) which are known to affect soil microbial function (Gopalakrishnan et al., 2007), and ratio of lignin syringyl to guaiacyl monomer units (Harman-Ware, Macaya-Sanz, et al., 2021; and personal communication with A. Harman-Ware) which is known to affect biomass recalcitrance to biochemical conversion (Alexander et al., 2020). Replicates of two of these selected genotypes (BESC-876 and BESC-119) had suffered mortality since the prior observations used for study design, so we were able to sample 69 individual live trees across those 24 genotypes.

Soil cores and roots were collected across two field campaigns in June and October 2022. Two adjacent soil cores were collected within 1 meter of the stem of each target tree using a 5 cm inner diameter slide hammer soil core sampler and split into increment depths of 0–15 cm and 15–30 cm (Fig. 2). The first set of cores was used for soil moisture, texture, and bulk density measurements, and the second set for fractionation and soil chemistry, as described in the next section. We also collected fine root samples from the surface (0–15 cm) layer for chemical analysis by tracing roots emerging from the target tree stems, through the root flare, to lateral roots, and then to fine roots bunches of diameter less than 2 mm. The collected soils and roots were stored in a cooler on blue ice (–4C) and shipped back to the lab for analysis.

**Figure 2.**
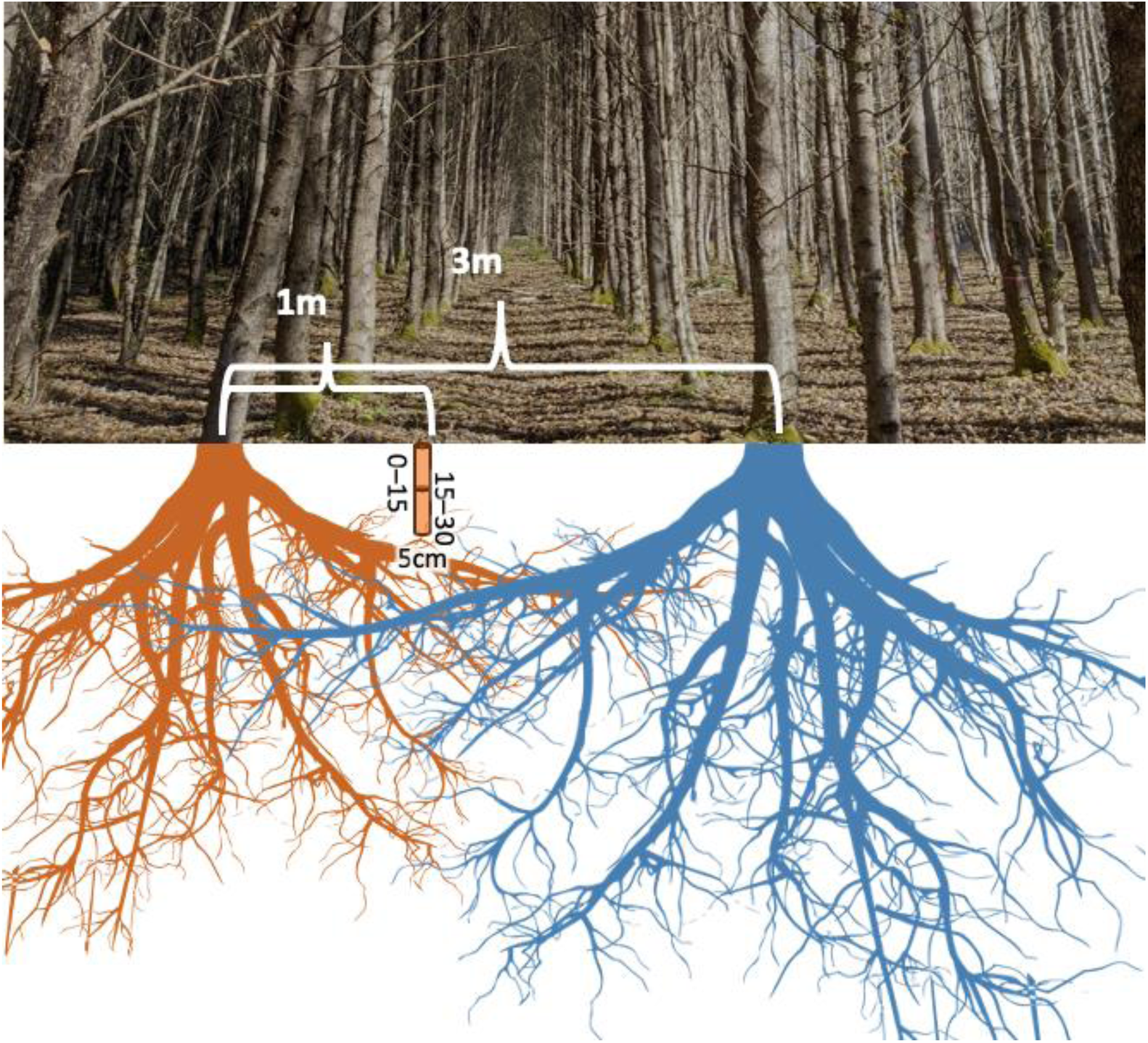
Schematic of poplar trial and soil sampling design, illustrating the location of soil core sampling relative to the rows of poplar trees and the overlap of their root systems. Scales are approximate (photo by the authors; root clipart from https://www.hiclipart.com/free-transparent-background-png-clipart-dvzrp).

Given the relatively tightly-space polyclonal grid planting of this site, we expected substantial root intermingling and enmeshing between neighboring trees which might obfuscate the soil carbon impacts of any individual tree. To quantify the amount of root intermingling in the zone from which we collected our soil cores, we also conducted exploratory root tracing on fine root bunches excavated randomly within 1 meter of 43 target trees. We found that 53% of those roots traced back to the target tree, 16% originated from a neighboring tree (in some cases more than one row away), and the remainder were of indeterminate origin (e.g., extended too deep for hand excavation). This confirmed that there is significant overlap of the root systems of these grid-planted trees (Fig. 2), however our soils were sampled in areas that likely contain a strong plurality of roots from the target tree.

### 2.3 Laboratory Analysis of Soils and Roots

Density core soil samples were evaluated for moisture, gravimetric water content, bulk density (dry soil mass divided by core volume; soil cores from this site were stone-free), and texture (Bouyoucos hydrometer method; Gee & Or, 2004), and the remaining sample mass was then air dried and archived. Soils from the chemistry cores were sieved to 2 mm and stored at 5°C until subsamples were taken and dried for further analysis. Soil pH was measured in a 0.005 M CaCl_2_ solution mixed 2:1 with soil using a glass electrode (Orion Versastar Advanced Electrochemistry Meter Thermo Scientific).

Additional subsamples from the 0–15 cm chemistry cores were separated into MAOM and POM fractions by size (Bradford et al., 2008; Cambardella & Elliott, 1992). A 5 g subsample of each soil was suspended in sodium hexametaphosphate (5% w/v) and placed on a reciprocal shaker (120 rpm) overnight to disperse aggregates. These solutions were then sieved (53 µm) with the aid of deionized water rinsing into a coarse POM fraction and a MAOM fine fraction. The recovered soil fractions were dried, weighed, ground, and analyzed for total carbon and nitrogen using an Elementar Unicube Trace CN analyzer (Elementar Corp., Ronkonkoma, NY).

Separate samples of bulk soil were air dried, ground, and analyzed for carbon and nitrogen in the same manner. In several cases the sum of the carbon recovered in the individual fractions differed from the carbon measured in the bulk samples by more than 30%, so these soils were re-fractionated. The final fractionation dataset had a median *c.* 17% under-estimate of total bulk SOC, likely attributable to MAOM losses during POM fraction rinsing or from particularly C-rich MOAM residues retained on the glassware after drying (J. Jastrow personal communication). We assume that carbonates were negligible in these acidic soils.

Dried and ground fine roots were sent to the University of Georgia Agricultural & Environmental Services Laboratories (AESL) for tissue chemistry analysis. Total aluminum (Al), boron (B), calcium (Ca), cadmium (Cd), chromium (Cr), copper (Cu), iron (Fe), potassium (K), magnesium (Mg), manganese (Mn), molybdenum (Mo), sodium (Na), nickel (Ni), phosphorus (P), lead (Pb), sulfur (S), and zinc (Zn) content were evaluated via microwave-assisted nitric acid digestion and inductively coupled plasma-mass spectrometry. Levels of root Cd, Mo, and Pb largely fell below instrument detection limits, so were excluded from further analysis. Cr was also omitted given it was not measured for a single genotype. Total carbon and nitrogen (N) were evaluated with an Elementar Vario Max Total Combustion Analyzer. Acid detergent lignin content was evaluated with an Ankom 2000 Fiber Analyzer to determine acid detergent fiber, followed by digestion in 72% sulfuric acid to isolate the lignin fraction. In addition, subsamples of bulk 0–15 cm soils were also sent to AESL for measurement of Mehlich-1 extractable nutrients (Ca, Cd, Cr, Cu, Fe, K, Mg, Mn, Mo, Na, Ni, P, Pb, Zn), cation exchange capacity (CEC), percent base saturation, and extractable Al. We excluded soil Cd, Cr, Mo, Pb from further analysis due to the lack of corresponding root concentration data.

### 2.4 Aboveground Trait Data

The Clatskanie common garden has been subject to extensive aboveground phenotyping since establishment, including periodic measurements of diameter at breast height (DBH; Chhetri et al., 2020; Evans et al., 2014; Slavov et al., 2024). We used an allometric equation from Truax et al. (2014) to estimate total aboveground biomass (stem + branch) from DBH (𝑏𝑖𝑜𝑚𝑎𝑠𝑠 = 0.071 · 𝐷𝐵𝐻^2.4055^) and estimated the annual productivity rate (i.e., growth) over the 10 years of observations (2009–2019). Note that for the 16 coppiced trees, we summed the aboveground biomass estimated from the DBH in 2012 (prior to coppicing) and the latest observed year, to estimate the total biomass growth over both coppice cycles.

### 2.5 Statistical Analysis

We used broad-sense heritability (𝐻^2^; Eqn. 1) to quantify the variance in soil C (including bulk soils and the individual POM and MAOM fractions) and above- and below-ground tree phenotypes attributable to tree genotype. For each phenotype and soil property, we estimated 𝐻^2^ by fitting a linear mixed effects model with genotype as a random effect (Chhetri et al., 2019) and soil texture, coppicing, and experimental block as fixed effects to control for spatial differences in environment and management, then extracted the estimated random effect 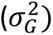 and residual variances 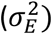. Additionally, we fit a mixed effects model with only a genotype random effect to check the importance of spatial fixed effects. All mixed-effects models were fit using the *lme4* package (Bates et al., 2015) in R version 4.4.1 (R Core Team, 2021).

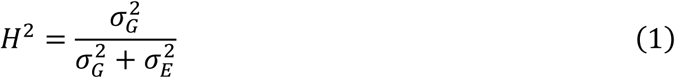

Next, we performed regularized regression to explore the effects of site environmental and management factors, soil properties, and tree traits on soil C in in the 0–15 cm surface layer.

Regularized regression reduces over-fitting and highly variable parameter estimates arising from correlated predictors (James et al., 2021). Specifically, we used the LASSO method in the *R* package *glmnet* (Friedman et al., 2010) within the *tidymodels* framework (Kuhn & Wickham, 2020), which also performs feature selection by shrinking less influential parameters to 0. We tuned the regularization parameter using 10-fold cross-validation repeated 10 times, then performed 10-fold cross-validation 20 times on the tuned model to estimate the sampling distributions of individual model parameters and overall model predictive performance. Using this procedure we tested three combinations of predictors to include in our model alongside block, soil texture, and coppicing: all root and soil chemistry predictors (“Root + Soil”), root chemistry only (“Root Only”), and soil chemistry only (“Soil Only”). Additional details and regularized regression results are given in Section S3. Our approach provides a reproducible analysis that objectively controls for correlated predictors and overfitting while quantifying the sensitivity of the regression to our data sample.

POM-C, MAOM-C, and total bulk SOC in the 0–15 cm layer were all individually analyzed for both broad-sense heritability and for associations with tree traits using the methods described above. Soil carbon concentration data (mg C/g dry soil as measured by the elemental analyzer) were analyzed directly. In addition, we analyzed per-area carbon stocks (calculated from carbon concentration and bulk density measurements) given their importance in quantifying the total carbon sequestration value of enhanced biosequestration in comparison to other natural climate solutions (Bossio et al., 2020; Paustian, Lehmann, et al., 2016). When calculating 𝐻^2^ for 0–15 cm total soil C stocks, we extracted the best-linear unbiased predictors of genotype level effects and used them to estimate the rate at which the carbon stocks under different genotypes have diverged over the 13 years since the common garden was first established.

All data and analysis code are currently available in a repository (https://github.com/sloan091/gcb-clatskanie-soil-c), and individual datasets will be published individually at https://cbi.ornl.gov/published-data/ at the time of manuscript publication.

Datasets include tree DBH and coppicing history from 2010–2019; fine root C, N, lignin, and elemental content from samples collected in 2022; and soil texture, pH, bulk density, and C & N content (both bulk and individual POM & MAOM fractions) from cores collected in 2022.

Genotype-level results for key soil properties and tree traits are summarized in Supplemental Table S1.

## 3 Results

### 3.1 SOC Concentrations and Stocks

Soil carbon measurements for the 0–15 cm soil cores collected under each of the 69 target trees are shown Figure 3, in both concentration and stock units. The median total SOC concentration is 52 mg C/g soil (ranges from 23–91 mg C/g soil), while the mineral-associated organic matter carbon (MAOM-C) and particulate organic matter carbon (POM-C) median values are 36 mg C/g soil (18–64 mg C/g soil) and 7 mg C/g soil (2–17 mg C/g soil), respectively. Associated median soil C/N ratios for bulk soil and the MAOM and POM fractions are 14 (9–16), 13 (10–15), and 20 (12–27), respectively. Combining the bulk soil C concentration data with bulk density, the median total 0–15 cm SOC stock at this site is approximately 68 tonnes/ha, comprised primarily of MAOM-C (48 tonnes/ha). The MAOM dominance is consistent with previous meta-analyses showing greater MAOM abundance under humid soil conditions (Heckman et al., 2023), managed landscapes (Lugato et al., 2021), and in temperate forests (Sokol et al., 2022). Mean SOC concentrations and stocks measured for individual genotypes are reported in Table S1.

**Figure 3.**
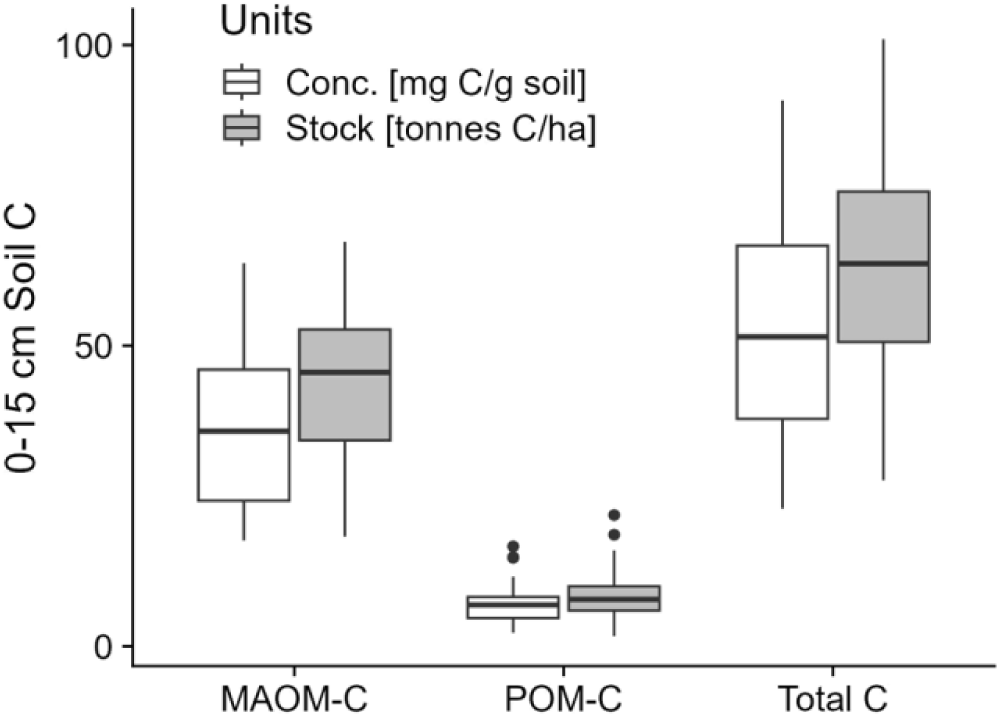
Soil carbon concentrations (conc.) and stocks measured for the mineral-associated organic matter (MAOM) fraction, particulate organic matter fraction (POM), and bulk soils (total C) collected from the 0–15 cm surface layer under 69 separate target trees in the Clatskanie common garden.

### 3.2 SOC and Tree Phenotype Heritability

Figure 4 shows the measured broad-sense heritability of SOC levels, aboveground biomass, and coupled root and soil chemistry observations. We observed no influence of tree genotype on total SOC concentration in the 0–15 cm surface layer, but individual MAOM-C and POM-C fractions showed marginal influence of genotype (𝐻^2^ ≈ 0.1). We found stronger influence of tree genotype on soil bulk density (𝐻^2^ = 0.23), and a strong inverse correlation between total SOC concentration and soil bulk density, which ranged from below 0.6 g/cm^3^ in high-SOC soils to above 1.0 g/cm^3^ in low-SOC soils (Fig. S3a). As a result, per-area 0–15 cm MAOM-C and POM- C stocks show over two times higher influence of tree genotype (𝐻^2^ ≈ 0.25) than their respective concentrations alone.

**Figure 4.**
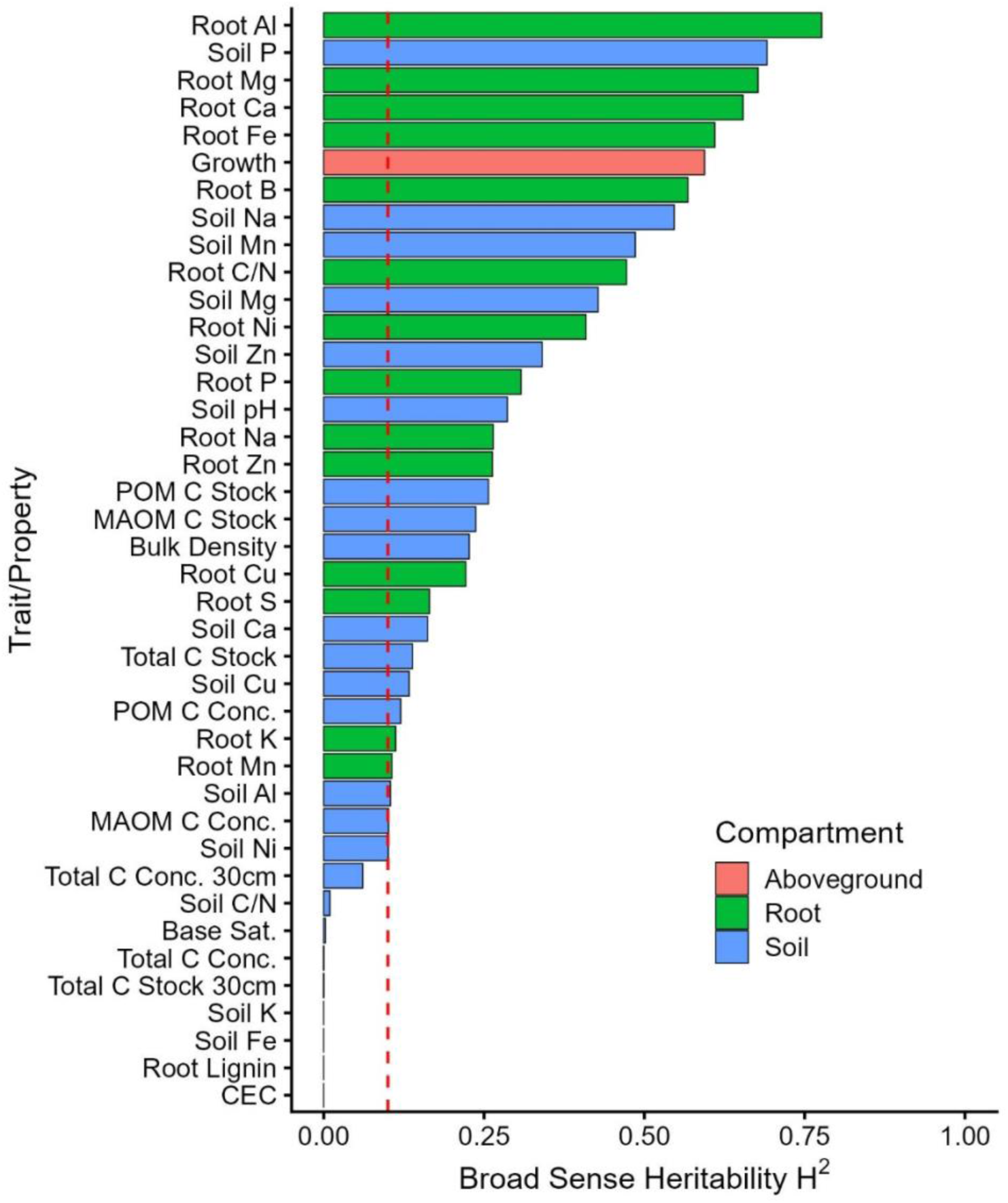
The broad sense heritability of various plant traits and soil properties across 24 genotypes at the Clatskanie common garden site. Biomass growth rate is the only aboveground property (red). Root properties (green) include the concentrations of various elements, plus lignin content and C/N ratio. Soil properties (blue) include the concentrations of various elements, soil pH, concentrations and stocks of particulate- and mineral-associated organic matter, bulk density, soil C/N ratio, base saturation (“Base Sat”), and cation exchange capacity (CEC). Soil properties are evaluated for the 0–15 cm surface layer unless otherwise noted. The red dashed line indicates 10% of the observed total variability is explained by genotype, a common threshold for evaluating heritable effects.

Both MAOM-C and POM-C are more influenced by tree genotype than is total SOC, regardless of whether stocks or concentration units are considered. The broad-sense heritability of MAOM-C is more sensitive than POM-C to background environmental variation (i.e., soil texture, coppicing and spatial block; Fig. S4), though this is consistent with longer-lived MAOM showing greater influence of pre-GWAS-establishment conditions than POM (Lavallee et al., 2020). We also collected 15–30 cm soil cores and analyzed for total SOC, but did not fractionate them. Interestingly, combining 0–15 and 15–30 cm samples into a composite 0–30 cm estimate reduced the apparent influence of tree genotype on both total SOC concentration (𝐻^2^ = 0.06) and stock (𝐻^2^ = 0). This might reflect a compounding of random measurement errors, and real inconsistencies in texture and other properties across the two soil layers (Fig. S3b). Soil pH appears moderately influenced by tree genotype, although it only varies from 5.2–5.6 across samples.

Plant productivity affects the input of biomass carbon available for stabilization to SOC. While we lack data on belowground carbon allocation or root biomass specifically, we found that the growth rate of aboveground biomass (genotype means ranging from 2.1–10.4 dry kg/tree/yr; Table S1) exhibited high heritability. Of the two phenotypes we measured as representative of root litter recalcitrance, root C/N ratio appears heritable (genotype means range from 46 to 92) and root lignin (28% to 37%) does not. The relatively low variability in root lignin across the 24 genotypes (Fig. S4c) may contribute to the lack of heritability; previous studies having identified a specific genetic locus controlling root lignin in hybrid poplar (Yin et al., 2010). Of the more than dozen types of root elements evaluated, many show significant variability (Fig. S4c) and high heritability (green in Fig. 4). Specifically, root Al, Fe, Ca, Mg, and B have more than 50% of their variability explained by genotype after controlling for site variables. Interestingly, bulk soil P, Na, Mn, and Mg also show high levels of tree genotype influence, and many other soil elements a moderate amount (blue in Fig. 4). There is generally little or no correlation between plant and soil element levels (Fig. S5), consistent with previous observations at this site (Craig et al., 2024). These results indicate that roots are not simply mirroring the surrounding soil chemistry and might even be selectively depleting certain elements from their surrounding soil.

### 3.3 Drivers of SOC Variability at Clatskanie

We next used regularized regression to quantify the effects of site factors, soil properties, and plant traits on the SOC at Clatskanie. We focus here on the MOAM-C concentration results because: 1) the MAOM-C concentration model had the best performance (median 𝑅^2^ ≈ 0.5; Section S3), 2) MAOM-C and Total C concentration and stock regressions had similar predictor effects, and 3) POM-C was poorly explained by the model given its small fraction at the site (Fig. 3). Additionally, we only analyze the regression results for all root and soil chemistry predictors (“Root + Soil”) and root chemistry predictors only (“Root Only”) as these two models had the best performance and many of the root and soil chemistry variables were heritable (Fig. 4).

Regardless, we provide all regression model results in Sect S3. Note, we omitted the foliar traits (ferulic acid, para-coumaric acid, and lignin S:G ratio) considered in the original genotype selection in this analysis given their lack of relationship with SOC (Fig. S6).

Figure 5 shows the potential drivers of MAOM-C at the Clatskanie site. Aboveground productivity factors (i.e., aboveground growth rate and coppicing, shown in red) generally have a small positive influence on MAOM-C concentration, approximately 1 mg C/g soil per standard deviation increase in the predictor. This is consistent with recent observations that MAOM-C is more influenced by net primary productivity than is POM-C (Hansen et al., 2024; King et al., 2023). The strong negative relationship (4–6 mg C/g soil per SD) between the finer soil texture (silt + clay) and MAOM-C runs counter to the expectation that MAOM-C is generally higher in soils with greater clay content that provides greater capacity for sorption to mineral surfaces (Hansen et al., 2024; Heckman et al., 2023). This may indicate that soil microbial activity and associated MAOM formation are lower in fine-textured, poorly-drained soils across this relatively wet site. The small negative effect of soil pH on MAOM (with “Roots Only” predictors) is possibly indicative of less microbial decomposition at lower pH, or of greater soil weathering at wetter, more acidic, sites leaving more secondary Fe and Al phases that effectively bind organics (Heckman et al., 2023). The latter mechanism is supported by the pH effect disappearing and a positive soil Al and Fe effect in the “Root + Soil” model.

**Figure 5.**
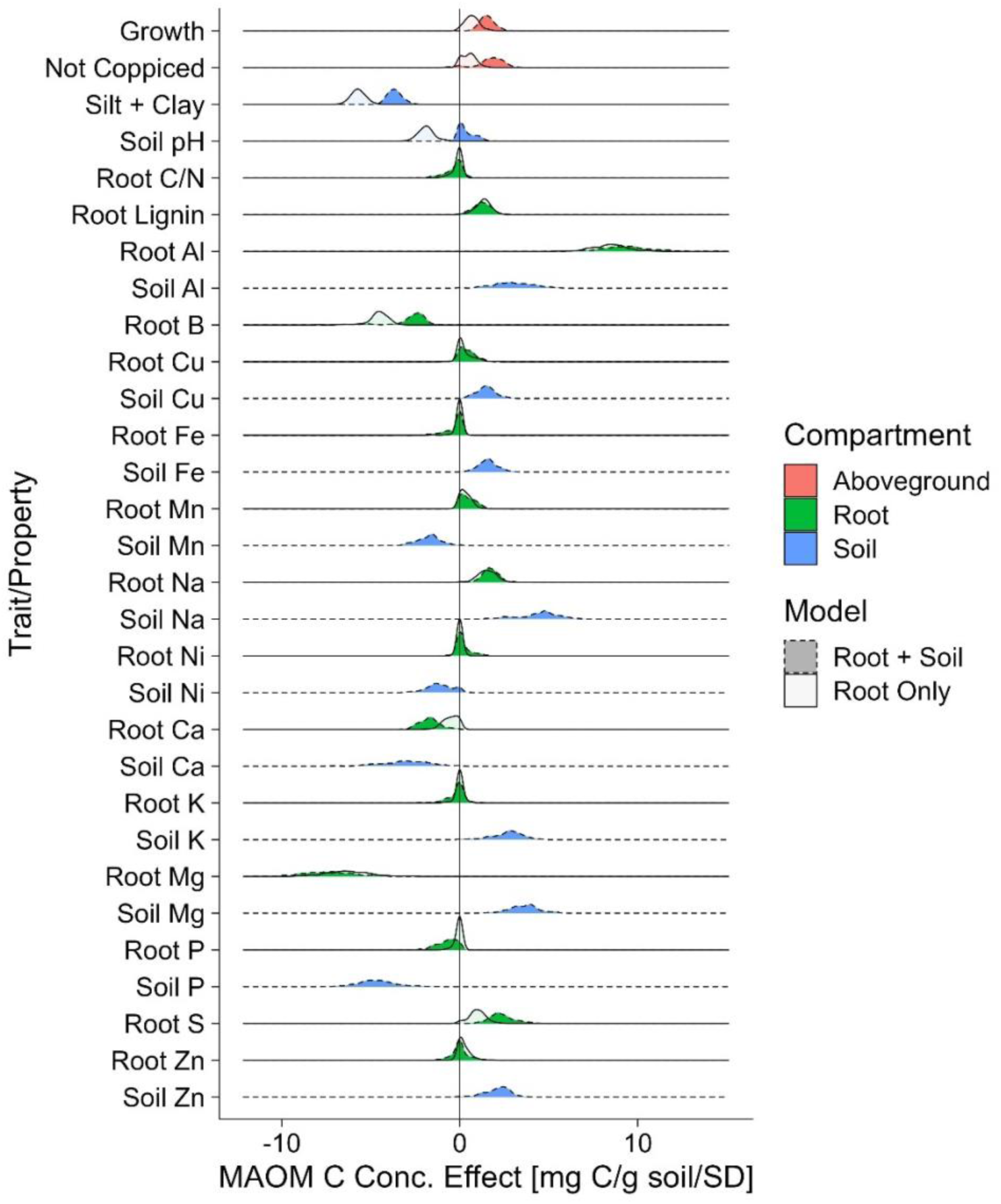
The effects of select aboveground, soil and root predictors on mineral associated organic matter (MAOM) C concentration. The predictor effects are standardized by their respective standard deviation (SD) and the distribution correspond to 200 repetitions of 10-fold cross-validation.

Regarding litter quality, root lignin has a moderate positive relationship with MAOM-C, counter to the notion in more modern SOC stabilization paradigms (Poirier et al., 2018) that lower-quality litter might increase POM at the expense of MAOM. This result, plus the positive soil Fe and Al effects, are consistent with previous work that has demonstrated preferential binding of lignin to iron and aluminum oxides (Hall et al., 2016; Heckman et al., 2013), which may be an important stabilization mechanism in the Wauna series soils that show redox mottling indicative of Fe oxide formation (National Cooperative Soil Survey, USDA, 2005). Finally, root C/N—an important SOC control in legacy biogeochemical models—has minimal effect on MAOM-C.

The largest effects on MAOM-C come from other aspects of root and soil chemistry, with root Al, Na, and S having a positive relationship with MAOM-C, and root Mg and B having a negative relationship. The positive root Al effects on MAOM-C match previous observations (Poirier et al., 2018), and could be due to the role of Al in precipitation of dissolved C (Scheel et al., 2008) or in reducing soil microbial activity (as has been previously observed in arbuscular mycorrhizal tree root decomposition; Zhao et al., 2023). Similarly, soil Al, Fe, Cu, Na, K, Mg, and Zn are positively associated with MAOM-C, while Mn, Ni, Ca, and P are negatively associated. The positive soil Al effect on MAOM-C is consistent with an inner-sphere bonding mechanism between organic matter and Al oxides (King et al., 2023; Rasmussen et al., 2018). The large opposite effects of root and soil Mg are noteworthy. The positive soil Mg effect is consistent with MAOM-C stabilization via cation bridging, yet we may not expect this mechanism to be as important at a wet, acidic site like Clatskanie given the relatively low Ca and Mg concentrations there (King et al., 2023).

### 3.4 Genotype Influence on Total Soil C Stock

We estimated the rate of divergence of SOC stocks under each genotype from the overall sample mean over the 13-year lifetime of the common garden, assuming the latter represents the initial SOC level across the garden in 2009. Figure 6 shows the genotype effect size on SOC stock estimated based on the mean of measured SOC stocks for each genotype (red) as reported in Table S1, and from the heritability analysis in Section 3.2 (which controls for soil texture, coppicing, and experimental block; teal). Genotypes with the highest and lowest SOC stocks are noted. The genotypes with the highest mean present-day SOC stocks (>80 tonnes C per ha) are BESC-35 (originally collected near Longview Junction, Washington, at the confluence of the Cowlitz and Columbia Rivers) and BESC-131 (originally collected from the banks of the Columbia River near Rainier, Oregon, approximately five miles further west). The genotype with the lowest mean SOC stock (30 tonnes C/ha) is GW-9768, which was originally collected from the South Fork of the Nooksack River near Acme, Washington.

**Figure 6.**
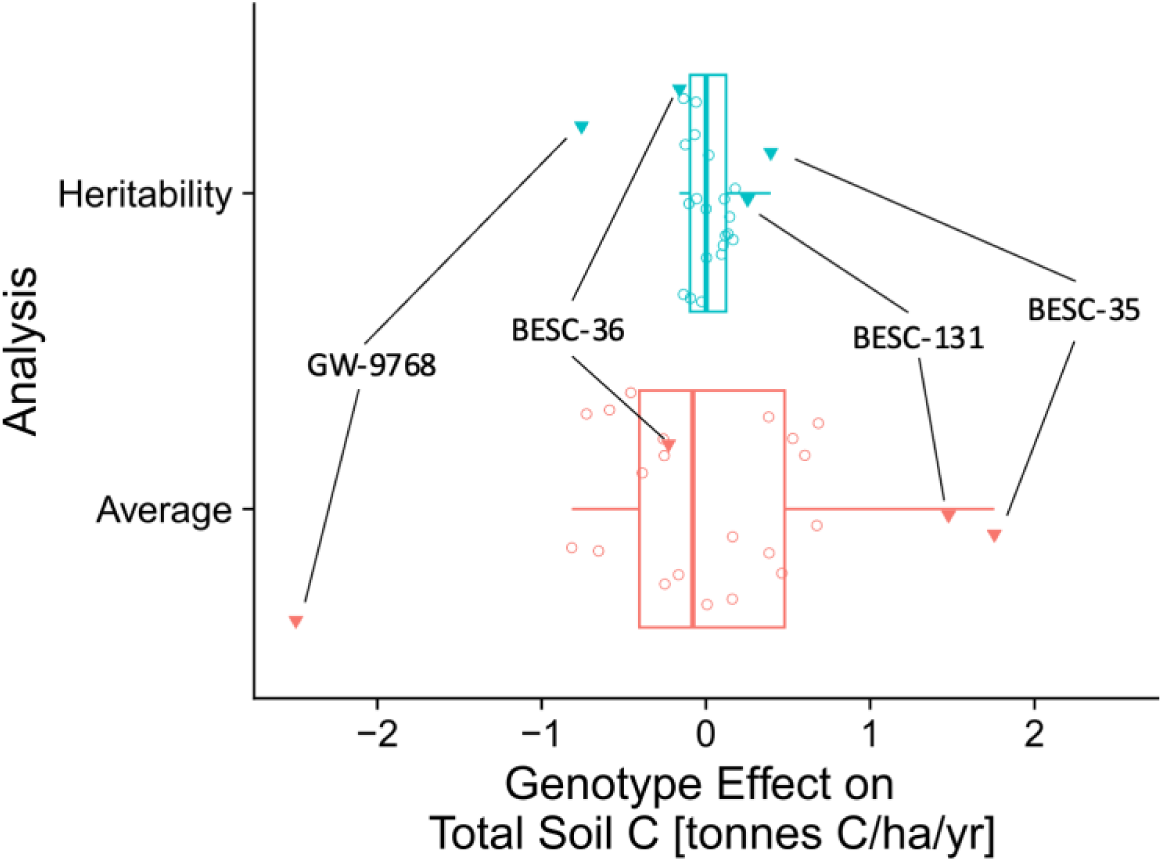
The estimated genotype effects on 0–15 cm total soil carbon stock change under the 24 genotypes at Clatskanie from 2009–2022. The analyses correspond to genotype estimates from the heritability analysis (teal) and a simple genotype average that does not account for the influence of site and management effects (red). The genotypes with the highest and lowest soil carbon effects as per the heritability analysis are noted.

The individual genotype means have diverged from the overall sample mean at rates between - 2.49 (GW-9768) to +1.76 tonnes C/ha/yr (BESC-35), with an interquartile range (IQR) of *c.* 0.8 tonnes C/ha/yr. The best linear unbiased predictors from the heritability analysis show that genotype effect ranges from -0.76 (GW-9768) to +0.39 tonnes C/ha/yr (BESC-35) with an IQR of *c.* 0.2 tonnes C/ha/yr. These heritability results reflect a conservative analysis shrunk towards the overall mean due to the small number of replicates and potentially large sampling uncertainty of soil core measurements. The true maximum range of SOC divergence between these genotypes likely lies between the mean and heritability estimates.

## 4 Discussion

Even with limited replication (n=3), we detected genotype-driven differences in MAOM-C stocks that account for ∼1/4 of the total observed MAOM-C stock variability (Fig. 4). Legacy SOC models like Century (Parton et al., 1987) and DayCent (Parton et al., 1998) consider plant productivity and litter quality as the main plant determinants of SOC. However, we found MAOM-C and total SOC stocks to be most strongly associated with root element concentrations (particularly Al, Mg, and B), with smaller effects from tree growth rate, root lignin, and other aspects of root chemistry (Fig. 5). These results are consistent with recent theory (Cotrufo et al., 2013) and experimental evidence (Ridgeway et al., 2022; Rossi et al., 2020) that increasing lignin content may be a relatively ineffective target for enhancing biosequestration. Instead, root elemental content was more strongly associated with MAOM-C levels, and was highly heritable across these genotypes, suggesting that these are plausible trait targets for plant breeding or engineering efforts. However, additional field observations and laboratory manipulation would be needed to validate these findings and pinpoint the underlying mechanism.

Together, these data provide a point of reference for the potential to enhance biosequestration by selecting for SOC-associated traits in *P. trichocarpa*. Even though we have only sampled a small portion of this GWAS population, the observed rates of SOC divergence between the highest-SOC genotype and the median suggest a potential for selection to improve sequestration by 0.3–1.7 tonnes C/ha/yr C in the 0–15 cm surface soil layer (Fig. 6). To put those rates in perspective, a meta-analysis by Qin et al. (2016) estimated that current commercial hybrid poplar grown on land previously under annual crops increases 0–30 cm soil C at an average rate of 0.3 tonnes C/ha/yr or loses soil C at a rate of 1.4 tonnes C/ha/yr when forest land is converted. Poplar field trials elsewhere in Oregon have shown no SOC differences compared to adjacent fields (Sartori et al., 2007), or sequestration of only 0.1 tonnes C/ha/yr (Collins et al., 2020).

The association between aboveground biomass and MAOM-C is potentially indicative of a trade-off between allocating additional carbon belowground for sequestration versus maintaining or increasing aboveground productivity (Jansson et al., 2021). While the total amount of carbon allocated belowground is known to be an important determinant of SOC, the limited association between aboveground biomass and MAOM-C present in this dataset implies that this trade-off may be less stark than previously thought. This may be because our aboveground biomass estimates fail to capture significant differences in allometry or litterfall dynamics between individual trees, because belowground allocation is highly variable in this population such that belowground inputs do not correlate with aboveground productivity, or because the physical or chemical form of those belowground inputs are more important than their magnitude.

### 4.1 Retroactively Evaluating SOC Changes in Common Gardens

Detecting any genotype-driven differences in soil carbon stocks in the Clatskanie common garden is notable since the site was not originally designed for biogeochemical research, and thus it lacked characterization of initial soil carbon at the time of establishment and is subject to significant root intermingling between neighboring trees in the polyclonal grid planting design. These results reinforce the value in measuring individual soil C fractions (Hansen et al., 2024; King et al., 2023; Lugato et al., 2021) rather than total SOC, which has been the focus of many previous common garden studies, but which did not show any influence of genotype at our site. Similarly, genotype effects were more evident on soil bulk density than soil carbon concentration, as observed in some previous studies (Craig et al., 2024; Waring et al., 2022). This study highlights the importance of measuring bulk density and evaluating C stocks (rather than just concentrations) of different organic matter fractions when attempting to detect SOC differences in common gardens.

Retrospective SOC analyses in common gardens are often made challenging due to limited biological replicates and statistical power, and a lack of initial SOC measurements at the time of establishment that could be subtracted from present day measurements to directly estimate SOC changes over time. In this analysis, we assumed that soil texture, coppicing, and spatial block capture the background environmental variability at Clatskanie. Although imperfect, these predictors would be static over the life of the garden and have a large influence on the heritability (Fig. S4) and regression results (Fig. 5). Our relatively disperse soil sampling across the common garden (Fig. 1B) precluded us from applying corrections to remove background environmental variability from our dataset, as has been done in some previous analyses of this population (Evans et al., 2014; Kristy et al., 2022). Future retroactive analyses of sites without initial SOC characterization should aim for denser soil sampling in order to enable more meaningful spatial corrections (Dupont et al., 2022).

Comparisons of carbon stocks based on repeated measurements of soil carbon concentration over time can also be confounded by corresponding changes in soil bulk density, such that different masses of soil are captured in fixed-depth cores collected at different points in time (von Haden et al., 2020). If a given management change or treatment simultaneously affects soil carbon concentration and bulk density—e.g., a change in tillage practice (Ogle et al., 2019) or grazing, or an application of an organic amendment (von Haden et al., 2020)—it can be appropriate to apply a correction to volumetric sampling results in order to compare carbon changes within a consistent mass of soil over time (Augusto & Boča, 2022; Ellert & Bettany, 1995; Lee et al., 2009). While we recognize the strong correlation between SOC and bulk density (Fig. S3a) implies that these trees have affected both carbon concentration and bulk density simultaneously, we have elected not to implement an equivalent soil mass correction in the interest of a more transparent and conservative analysis and for consistency with previous studies (e.g., Waring et al., 2022).

### 4.2 Effects of Root Elemental Content on MAOM-C Stabilization

The outsized effects of root and soil nutrients and trace elements suggest those might be better targets for plant bio-design than lignin content or C/N ratio, at least with respect to this site.

Low leaf Mg (Yang et al., 2022), low fine root Ca (Silver & Miya, 2001), and high fine root Al (in arbuscular mycorrhizal-associated trees; Zhao et al., 2023) have all been associated with higher recalcitrance (i.e., lower short-term litter decomposition rates) of those tissues. In our dataset, these same properties in fine roots are associated with higher MAOM-C levels. However, root effects on mineral association are less well understood than their effects on decomposition and soil aggregation (Poirier et al., 2018), and root elemental content does not necessarily cleanly map to older versus more modern SOC stabilization paradigms. The large positive effects of both root Al and soil Al on MAOM-C observed here match previous results in acidic, wet forests (Hobbie et al., 2007; Poirier et al., 2018; Russell et al., 2007; Scheel et al., 2008), though potentially through differing mechanisms. Scheel et al. (2008) performed soil incubation experiments to show that root Al helps precipitate dissolved C, slowing further microbial decomposition via reduced bioavailability rather than Al toxicity to microbes. The negative effects of soil P on MAOM-C shown here also match those incubation results. Alternately, the positive effect of soil Al and Fe on MAOM-C matches stabilization via an inner-sphere bonding mechanism common in temperate forests, and their inclusion in the model causes the negative soil pH effect on MAOM-C to disappear (Fig. 5; Heckman et al., 2023; King et al., 2023; Rasmussen et al., 2018).

However, when evaluating these results, it is important to note that the tight correlations between some elements (e.g., Al, B, and Mg; Fig. S7) makes their individual effects difficult to discern, even using regularized regression. Furthermore, heritability analysis (Fig. 4) suggests that there is a tree genotype effect on levels of certain elements in the surrounding soils, presumably due to plant uptake and depletion, or transfer of elements from deep soils to surface soil horizons over time (Oostra et al., 2006; Reich et al., 2005). To the extent that these trees are selectively enhancing or depleting specific elements in the surface soil, it is even more challenging to separate those effects from the effects of root concentrations of those same elements on MAOM-C. The lack of initial soil characterization at the time of establishment makes it difficult to interpret these results, though future campaigns to sample root-associated rhizosphere soils specifically could be helpful.

### 4.3 Limitations and Future Work

It is important not to over-generalize the results from this single site. The Clatskanie common garden is established in a marginal lowland site with poorly drained alluvial soils and with a shallow water table and prone to flooding. A drier site might show different soil texture and chemistry effects or greater presence of POM-C (Heckman et al., 2023). Given the outsized importance of MAOM-C at Clatskanie, future belowground sampling efforts should also consider root exudation, an under-studied plant trait and possible biosequestration target (Jansson et al., 2021; Panchal et al., 2022). Root exudates provide a source of carbon for microbial biomass growth, a source of organic compounds that can be directly sorbed to soil minerals, or in the case of organic acids, a priming agent that de-stabilizes existing MAOM (Clarholm et al., 2015; Keiluweit et al., 2015). Thus, it is important to quantify both the magnitude of root exudation and its chemical quality in order to understand whether it will have a net positive or negative effect on SOC (Poirier et al., 2018).

Finally, our low spatial sampling density and low number of available biological replicates per genotype (n=3) limits the statistical power of our analysis of individual genotype SOC stock change rates (Fig. 6), leaving those results susceptible to error (Gelman & Carlin, 2014) and with limited potential for spatiotemporal extrapolation. However, belowground sampling under a larger portion of this GWAS population might allow us to link any observed differences with specific single nucleotide polymorphisms and alleles across these sequenced genotypes with much greater statistical precision. Belowground sampling of roots and especially soil carbon is typically laborious and expensive (Paustian, Collier, et al., 2019). To the extent that studies like this can inform what types of soil C measurements are most likely to show genotype influences, and which belowground plant traits are most strongly associated with SOC outcomes, it potentially paves the way for more targeted sampling campaigns at GWAS scales that might elucidate the biological controls on SOC with greater detail and certainty.

## 5 CRediT author statement

JLF: Conceptualization, Formal analysis, Funding acquisition, Investigation, Methodology, Project administration, Visualization, Writing – original draft, review & editing.

BPS: Conceptualization, Data Curation, Formal analysis, Investigation, Methodology, Visualization, Writing – original draft, review & editing.

MEC: Conceptualization, Formal analysis, Funding acquisition, Investigation, Methodology, Writing – review & editing.

PC: Investigation, Writing – review & editing.

SLO: Investigation, Methodology, Writing – review & editing. TM: Investigation, Writing – review & editing.

RZA: Conceptualization, Funding acquisition, Investigation, Methodology, Writing – review & editing.

MPV: Data Curation, Writing – review & editing.

HBC: Investigation, Methodology, Writing – review & editing.

KH: Investigation, Resources.

UCK: Supervision, Writing – review & editing. WM: Supervision, Writing – review & editing.

CWS: Conceptualization, Investigation, Methodology, Supervision, Writing – review & editing. MAM: Conceptualization, Funding acquisition, Investigation, Methodology, Project administration, Supervision, Writing – original draft, review & editing.

## Acknowledgements

This work was supported through Oak Ridge National Laboratory (ORNL) Laboratory Directed Research and Development (LDRD) Projects 11146, 10681, and 10978. CWS was supported by the Bio-Scales project under the Genomic Sciences Program, U.S. Department of Energy, Office of Biological and Environmental Research (BER) under contract number DE-AC05-00OR22725. This material is based upon work supported by the Center for Bioenergy Innovation (CBI), U.S. Department of Energy, Office of Science, Biological and Environmental Research Program under Award Number ERKP886. We thank Rick Stonex and Brian Stanton of Poplar Innovations for hosting, advice, and assistance around fieldwork; Geoff Schwaner for help with soil sample elemental analysis; Stanton Martin for help accessing previously collected aboveground phenotype data; Chanaka Roshan Abeyratne for feedback on spatial analysis methods; and Anne “Liz” Harman-Ware and Timothy J. Tschaplinski for their help accessing and interpreting foliar lignin and metabolomic data, respectively. We also thank Elizabeth Herndon and Francesca Cotrufo for their helpful reviews of earlier versions of this manuscript. ORNL is managed by UT-Battelle, LLC for the U.S. DOE under contract DE-AC05-00OR22725.

## 6 Data availability statement

All data and code associated with this manuscript will be made publicly accessible with a permanent DOI through the Center for Bioenergy data repository (https://cbi.ornl.gov/published-data/) at the time of publication, but currently reside at https://github.com/sloan091/gcb-clatskanie-soil-c. Datasets include tree DBH and coppicing history from 2010–2019; fine root C, N, lignin, and elemental content from samples collected in 2022; and soil texture, pH, bulk density, C & N content (both bulk and individual POM & MAOM fractions) and plant-available nutrient data from cores collected in 2022.

