## Supplemental Information for "Which plant traits increase soil carbon sequestration? Empirical evidence from a long-term poplar genetic diversity trial"

**Authors:** John L. Field[1,*] (https://orcid.org/0000-0003-4451-8947), Brandon P. Sloan[1] (https://orcid.org/0000-0003-1630-4271), Matthew E. Craig[1] (https://orcid.org/0000-0002-8890-7920), Parker Calloway[2] (https://orcid.org/0000-0002-9269-2993), Sarah L. Ottinger[3] (https://orcid.org/0000-0002-3011-8523), Thomas Mead[1] (https://orcid.org/0009-0009-0415-5329), Rose Z. Abramoff[4,5] (https://orcid.org/0000-0002-3393-3064), Mirko Pavicic Venegas[6] (https://orcid.org/0000-0002-2908-3183), Hari B. Chhetri[6] (https://orcid.org/0000-0001-6820-8789), Kathy Haiby[7], Udaya C. Kalluri[6] (https://orcid.org/0000-0002-5963-8370), Wellington Muchero[6] (https://orcid.org/0000-0002-0200-9856), Christopher W. Schadt[6] (https://orcid.org/0000-0001-8759-2448), Melanie A. Mayes[1] (https://orcid.org/0000-0001-6368-9210)

*

[1]Environmental Sciences Division, Oak Ridge National Laboratory, Oak Ridge, TN

[2]Dept. of Civil, Construction, and Environmental Engineering, North Carolina State University, Raleigh, NC

[3]Tennessee Valley Authority, Knoxville, TN

[4]School of Forest Resources, University of Maine, Orono, ME

[5]Wintergreen Earth Science, Kennebunk, ME

[6]Biosciences Division, Oak Ridge National Laboratory, Oak Ridge, TN

[7]Poplar Innovations, Inc., Castle Rock, WA

*Notice: This manuscript has been authored by UT-Battelle, LLC, under contract DE-AC05-00OR22725 with the US Department of Energy (DOE). The US government retains and the publisher, by accepting the work for publication, acknowledges that the US government retains a non-exclusive, paid-up, irrevocable, world-wide license to publish or reproduce the submitted manuscript version of this work, or allow others to do so, for US government purposes. DOE will provide public access to these results of federally sponsored research in accordance with the DOE Public Access Plan (https://energy.gov/doe-public-access-plan).*

### 1 Supplementary Materials

**S1. Supplemental Tables**

*Table S1. Select genotype-mean (standard deviation) soil and root properties. Full data & code available at https://cbi.ornl.gov/published-data/.*

| Genotype | Soil pH | Soil bulk density ($g/cm^3$) | POM-C (mg C/g soil) | MAOM-C (mg C/g soil) | Total soil C (mg C/g soil) | Total soil C stock (Mg C/ha) | Aboveground biomass (kg/y/tree) | Root C/N | Root lignin (%) | Root Al (%) | Root Mg (%) | Root B (ppm) |
|---|---|---|---|---|---|---|---|---|---|---|---|---|
| BESC-102 | 5.34 (0.12) | 0.76 (0.13) | 9.51 (1.99) | 48.2 (7.5) | 63.4 (11.7) | 71.5 (8.3) | 5.76 (2.11) | 46.7 (5.9) | 27.8 (4.5) | 0.69 (0.17) | 0.16 (0.03) | 15.0 (2.9) |
| BESC-119 | 4.78 (nan) | 0.82 (nan) | 7.07 (nan) | 32.0 (nan) | 42.4 (nan) | 52.0 (nan) | 3.34 (nan) | 51.9 (nan) | 35.7 (nan) | 0.21 (nan) | 0.1 (nan) | 11.8 (nan) |
| BESC-13 | 5.28 (0.04) | 0.93 (0.08) | 8.41 (6.02) | 26.1 (4.0) | **39.3 (12.9)** | 54.1 (15.1) | 4.11 (2.56) | **46.2 (5.7)** | 29.7 (2.4) | 0.17 (0.06) | 0.11 (0.01) | 14.9 (0.5) |
| BESC-131 | 4.94 (0.14) | 0.78 (0.08) | 7.45 (1.94) | 50.9 (3.6) | 70.0 (4.8) | **81.8 (2.9)** | 4.83 (2.41) | 61.7 (11.8) | 33.1 (6.5) | 0.29 (0.08) | 0.09 (0.02) | 10.8 (0.8) |
| BESC-133 | 5.1 (0.19) | 0.68 (0.12) | 5.09 (2.4) | 39.5 (20.5) | 56.9 (30.5) | 55.0 (23.8) | 8.56 (3.89) | 56.0 (13.1) | 35.0 (3.0) | 0.24 (0.04) | 0.11 (0.02) | 11.2 (2.0) |
| BESC-144 | 5.37 (0.17) | 0.84 (0.03) | 7.08 (1.26) | 40.7 (3.5) | 53.4 (3.2) | 67.6 (6.2) | 8.6 (7.26) | 67.7 (9.9) | 30.4 (2.8) | 0.25 (0.06) | 0.1 (0.02) | 11.7 (1.8) |
| BESC-145 | 5.33 (0.24) | 0.89 (0.25) | 8.48 (2.03) | 34.9 (17.1) | 52.8 (23.5) | 64.7 (10.4) | 7.53 (1.61) | **91.7 (13.8)** | 29.2 (8.3) | 0.23 (0.04) | 0.12 (0.01) | 11.9 (1.0) |
| BESC-192 | 5.0 (0.3) | 0.91 (0.28) | 5.52 (2.06) | 34.1 (25.7) | 50.0 (35.4) | 59.2 (22.9) | 9.69 (1.57) | 64.0 (8.9) | 33.3 (5.5) | 0.26 (0.07) | 0.11 (0.03) | 10.7 (1.1) |
| BESC-2 | 5.18 (0.27) | 0.82 (0.15) | 5.48 (1.73) | 34.4 (22.6) | 49.5 (34.6) | 56.7 (29.8) | 5.5 (0.63) | 57.0 (6.7) | 29.3 (7.8) | 0.21 (0.06) | 0.1 (0.02) | 10.5 (2.2) |
| BESC-265 | 5.06 (0.2) | 0.64 (0.05) | 10.64 (3.88) | 51.8 (1.4) | **75.0 (1.6)** | 71.4 (4.5) | 7.54 (1.35) | 50.2 (11.9) | 33.4 (1.7) | 0.28 (0.02) | 0.11 (0.01) | 10.7 (1.3) |
| BESC-319 | 5.12 (0.27) | 0.9 (0.2) | 8.76 (2.68) | 34.4 (16.6) | 54.9 (20.3) | 70.4 (10.4) | 8.86 (3.2) | 57.6 (4.7) | 31.8 (2.9) | 0.27 (0.04) | 0.11 (0.02) | 11.0 (1.4) |
| BESC-35 | 4.99 (0.06) | 0.95 (0.07) | 11.22 (5.37) | 38.0 (11.1) | 61.1 (20.4) | **85.4 (24.8)** | 5.17 (1.63) | 63.2 (14.1) | **27.6 (2.3)** | 0.43 (0.21) | 0.13 (0.03) | 12.3 (2.6) |
| BESC-351 | 5.06 (0.12) | 0.89 (0.12) | 6.24 (1.92) | 35.4 (7.6) | 51.4 (8.9) | 67.6 (7.2) | 5.65 (1.29) | 53.4 (4.0) | 31.2 (2.5) | 0.29 (0.03) | 0.1 (0.02) | 9.5 (1.1) |
| BESC-36 | 5.1 (0.17) | 0.83 (0.1) | 5.39 (1.82) | 37.1 (14.9) | 49.6 (20.7) | **59.7 (17.2)** | 2.27 (0.84) | 54.7 (4.4) | 32.4 (1.9) | 0.26 (0.12) | 0.1 (0.03) | 11.3 (1.4) |
| BESC-388 | 5.1 (0.34) | 0.86 (0.12) | 5.36 (1.92) | 35.8 (20.2) | 47.9 (33.6) | 59.4 (35.3) | 3.34 (2.01) | 50.8 (2.6) | **36.4 (9.4)** | 0.36 (0.1) | 0.11 (0.02) | 11.6 (1.8) |
| BESC-833 | 5.1 (0.11) | 0.77 (0.07) | 6.33 (1.92) | 36.4 (9.6) | 50.6 (11.3) | 59.3 (18.0) | 8.51 (2.62) | 73.6 (11.6) | 30.1 (3.4) | 0.2 (0.09) | 0.09 (0.01) | 9.4 (1.3) |
| BESC-841 | 5.05 (0.07) | 0.91 (0.01) | 4.53 (0.65) | 30.9 (6.6) | 45.8 (9.2) | 62.7 (12.1) | 5.67 (0.72) | 60.6 (10.3) | 35.9 (4.1) | 0.26 (0.05) | 0.1 (0.01) | 11.2 (1.0) |
| BESC-876 | 5.2 (0.26) | 0.94 (0.06) | 6.77 (1.35) | 33.8 (5.2) | 49.3 (14.0) | 68.6 (15.6) | **10.35 (3.31)** | 59.9 (11.4) | 28.7 (4.3) | 0.51 (0.03) | 0.15 (0.0) | 15.1 (1.8) |
| BESC-897 | 5.18 (0.17) | 0.83 (0.11) | 6.27 (1.53) | 33.9 (7.0) | 50.9 (11.6) | 64.7 (23.0) | 7.88 (3.29) | 71.9 (6.1) | 35.6 (4.3) | 0.18 (0.05) | 0.09 (0.01) | 9.9 (1.0) |
| GW-11026 | 5.13 (0.23) | 0.83 (0.11) | 5.06 (1.53) | 35.9 (11.4) | 49.6 (13.8) | 60.4 (11.2) | 2.82 (0.72) | 61.4 (7.1) | 31.2 (6.8) | 0.4 (0.05) | 0.14 (0.02) | 14.3 (2.4) |
| GW-9768 | 5.19 (0.14) | 0.57 (0.25) | 4.8 (3.23) | 26.7 (10.6) | **39.3 (14.5)** | **30.2 (2.5)** | 6.27 (1.76) | 76.2 (15.9) | 29.4 (6.0) | 0.26 (0.13) | 0.13 (0.02) | 13.6 (2.1) |
| GW-9776 | 5.39 (0.4) | 0.86 (0.21) | 5.75 (4.29) | 34.8 (20.6) | 49.5 (33.5) | 57.6 (26.5) | 8.53 (1.1) | 65.7 (5.1) | 29.7 (1.5) | 0.56 (0.2) | 0.16 (0.03) | 19.3 (7.3) |
| GW-9854 | 5.3 (0.12) | 0.88 (0.14) | 5.67 (2.75) | 28.7 (12.3) | 42.0 (19.3) | 53.2 (15.5) | **2.07 (0.24)** | 66.7 (17.1) | 31.4 (14.1) | 0.22 (0.05) | 0.1 (0.01) | 11.7 (0.7) |
| SLMC-28-2 | 5.01 (0.11) | 0.8 (0.13) | 7.8 (2.32) | 38.9 (8.0) | 59.0 (12.4) | 69.5 (9.3) | 2.49 (0.65) | 62.8 (7.6) | 29.3 (5.7) | 0.29 (0.12) | 0.11 (0.02) | 11.8 (2.8) |
| **Population** | **5.15 (0.21)** | **0.83 (0.15)** | **6.86 (2.94)** | **36.6 (12.8)** | **52.6 (18.6)** | **62.8 (18.0)** | **6.07 (3.18)** | **61.6 (13.1)** | **31.5 (5.32)** | **0.31 (0.15)** | **12.1 (2.9)** | **0.11 (0.03)** |

#### S2. Supplemental Figures

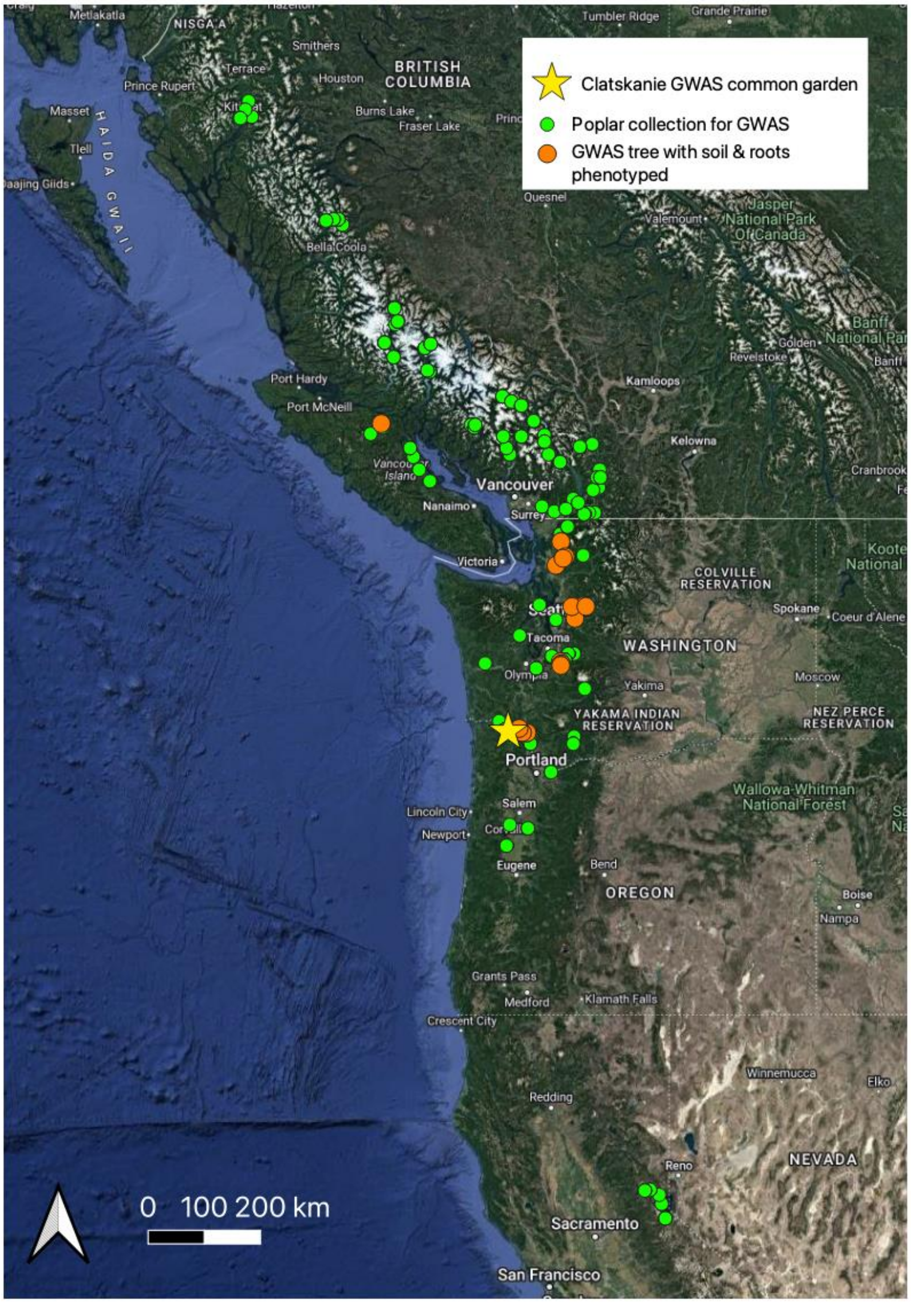


*Figure S1. Origin of black cottonwood (Populus trichocarpa) trees included in the GWAS population, highlighting those trees that were phenotyped for belowground traits and soil carbon outcomes in the current study.*

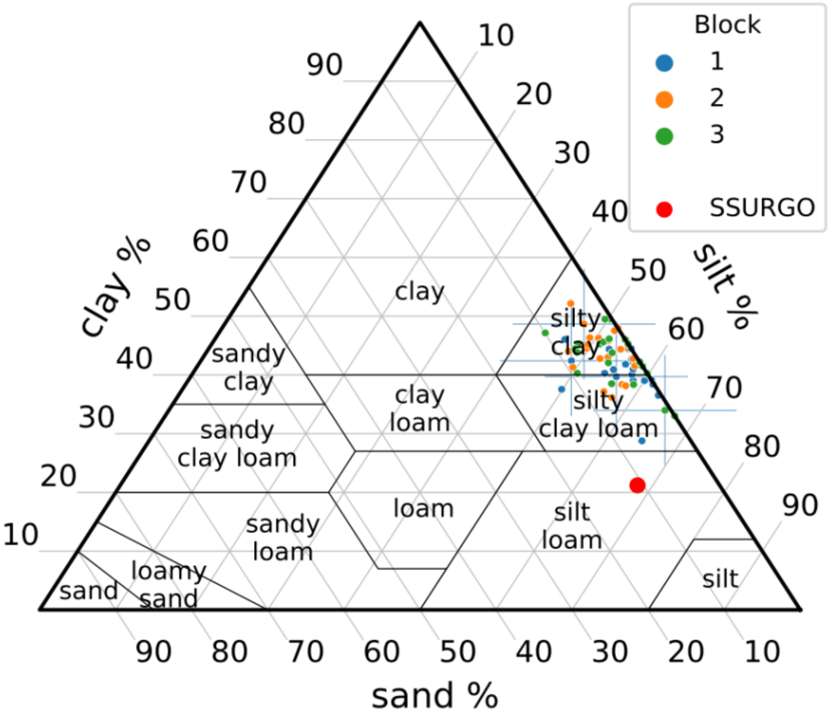


*Figure S2. Measured 0–30 cm soil texture data displayed on a soil texture triangle. The texture of the Wauna silt loam expected at this as per the SSURGO database is shown in red.*

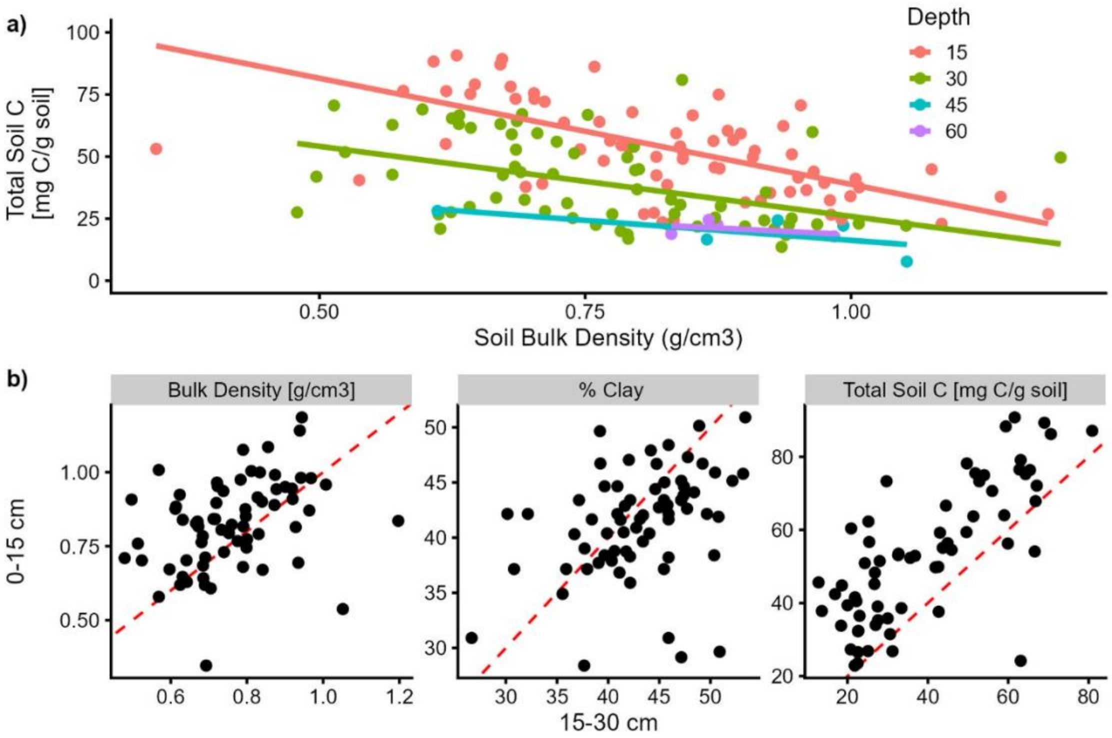


*Figure S3. Comparison of differing soil core depth observations. A) Total soil C concentration and bulk density are inversely related but with gradually decreasing slopes at deeper depths. B) Relationship between soil bulk density, texture (clay content), and total SOC concentration measured at each tree in the 0–15 versus the 15–30 cm soil layer. Although correlated, there is much variability between measures at differing depths that likely confounds heritability analysis when combining data from both depths into a single 0–30 cm value.*

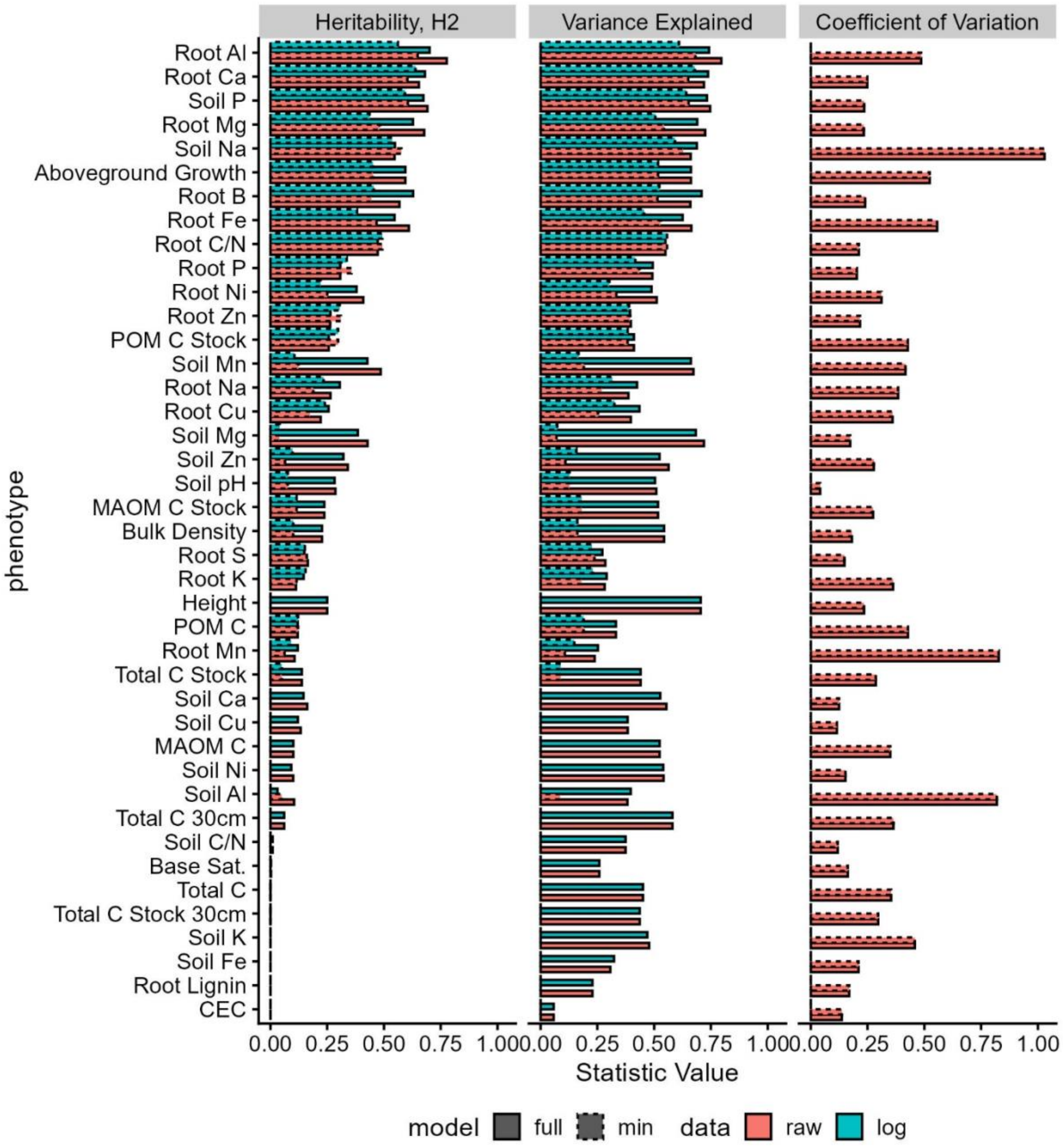


*Figure S4. The heritability results from fitting the linear mixed effects model with ("full") and without ("min") the background covariates of soil texture, coppicing and block as well as with the phenotype log-transformed. Here we show the a) broad-sense heritability ($H^2$), b) the variance explained by the model, and c) the coefficient of variation (standard deviation/mean) for each phenotype. Note that we only show the coefficient of variation for the raw data, as the statistic is not meaningful in the log-space.*

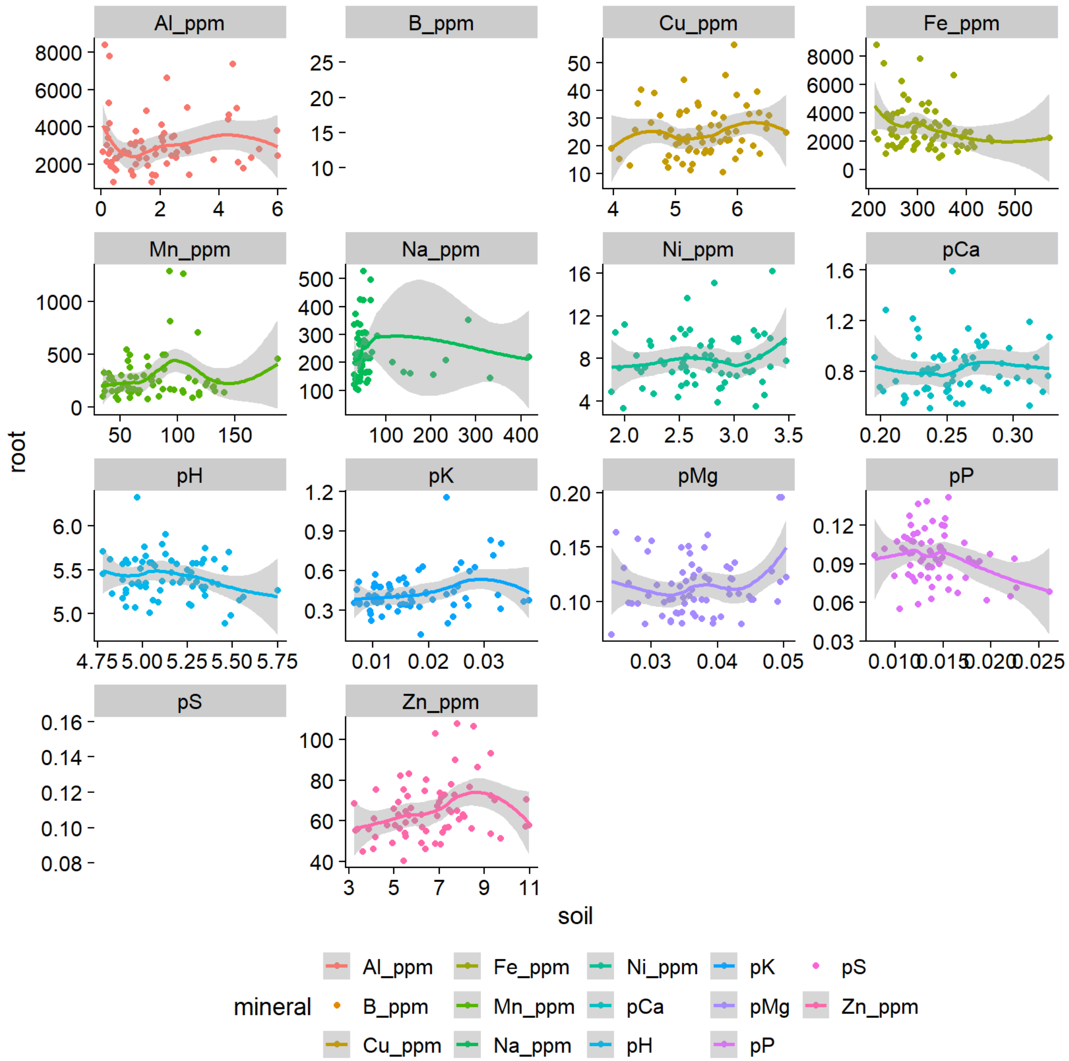


*Figure S5. The relationship between root and soil elemental content for the 69 trees at Clatskanie. Note that there are not typically strong relationships between the two. Units are noted in the subplot titles as either percent ("p") or parts per million ("ppm").*

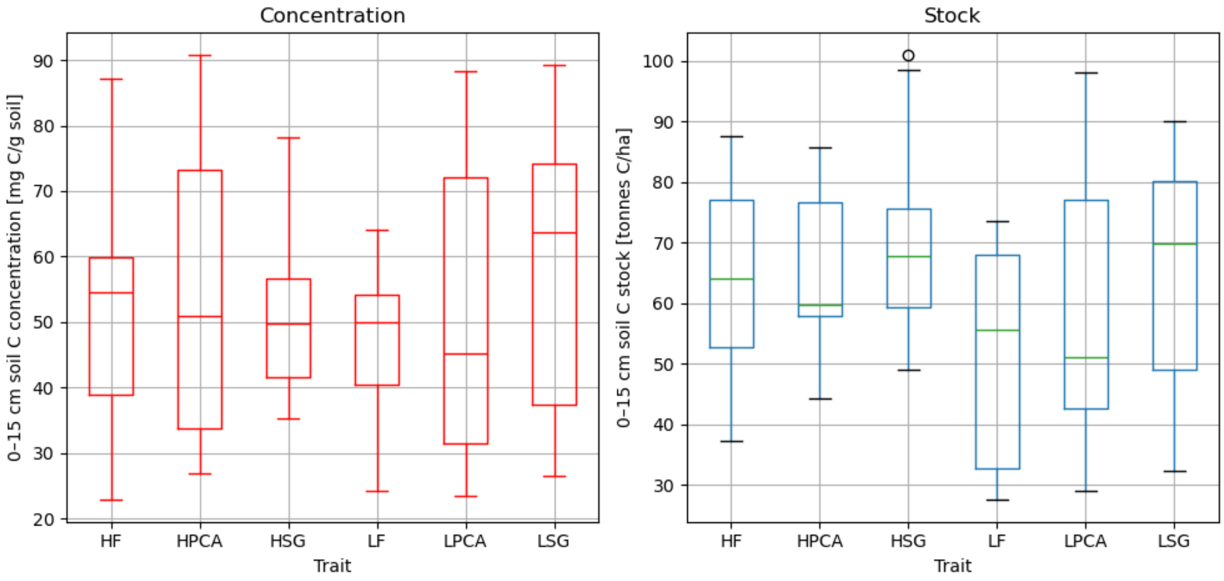


*Figure S6. Total soil carbon concentrations and stocks for trees grouped by traits used in initial experimental design. Genotypes were selected for sampling and analysis for having **h**igh or **l**ow values of foliar ferulic acid (HF and LF), para-coumaric acid (HPCA and LPCA), and lignin S:G ratio (HSG and LSG).*

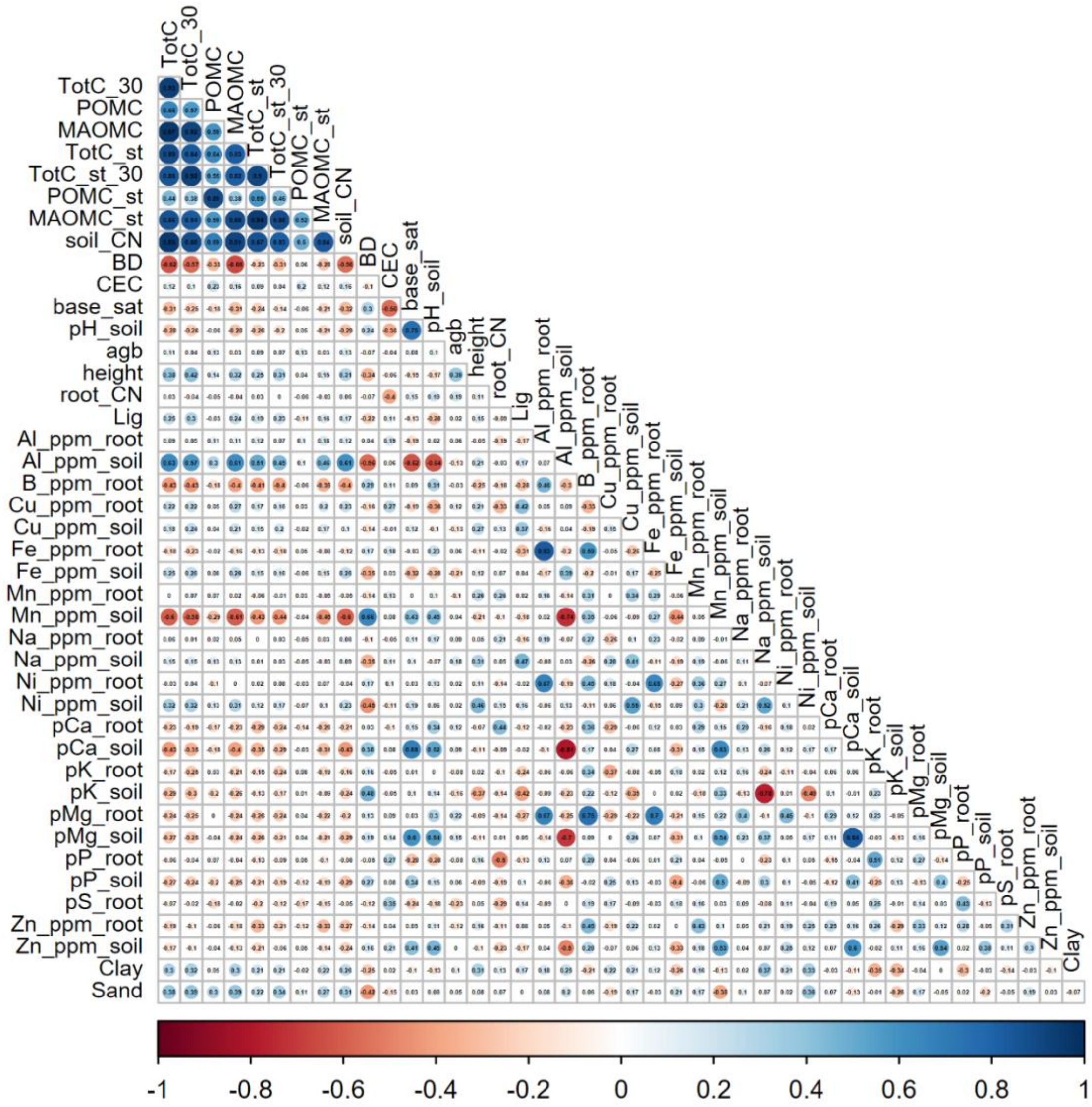


*Figure S7. Correlation matrix showing relationship between various measured root and soil nutrients and trace elements, soil texture, and pH.*

**S3. Further Details on Regularized Regression**

We fit linear models for three sets of predictors with and without log-transformation for the MAOM, POM and total C concentrations and stocks using two different algorithms; these various permutations result in 72 differing models. The three predictor sets all contained site, soil texture, and aboveground growth predictors, but differed in whether they include root nutrients and trace elements only (“root”), soil nutrients and trace elements only (“soil”), or both sets (“all”). Section 3.3 focuses on the “all” and “root” scenarios, since root nutrients were found to be critical determinants of MAOM-C and total C.

The large number of correlated predictors could result in model over-fitting and highly variable effects. To combat this, we performed regularized regression and cross-validate the results to get a better estimate of the uncertainty in effects and model predictive performance. We performed both ridge and LASSO regression using the R package *glmnet* contained in the *tidymodels* ecosystem. The two regularized regression methods are end-members of elastic-net regression, which penalizes parameter norms causing a shrinkage in parameter values (some to 0 in the case of LASSO) that biases the model fit to the training sample while improving the model's ability to fit to out-of-bag samples (Hastie et al., 2009)[1]. We performed our regression in two steps: 1) tune or train the model to determine the penalty hyperparameter using 10-fold cross-validation repeated 10 times, and 2) fit the model with the optimal penalty parameter to

[1] Hastie, T., Tibshirani, R., & Friedman, J. (2009). The Elements of Statistical Learning. Springer. https://doi.org/10.1007/978-0-387-84858-7

random samples of the data (via repeated cross-validation) to get at the sampling distribution of the effects and predictive performance.

The results of the hyperparameter tuning are shown in Figure S8 for the “root” predictor model only (for brevity). We defined the optimal penalty parameter (vertical lines) as the value with the lowest root mean square error (RMSE) on the held-out test set in the cross-validation. We note here that MAOM and Total C have near identical results, reflecting the dominance of the MAOM-C fraction at Clatskanie (Fig. 3). Additionally, the POM-C models do not adequately fit the data—the decay in performance with increasing penalty and normalized RMSE values greater than 1 together indicate that the mean is a better predictor than our model. The model also appears to fit better to concentration rather than stock units, and to raw data rather than log-transformed data. Lastly, the LASSO method appears to provide a small predictive advantage over the ridge regression. For all of these reasons, we focused on MAOM-C concentration results using raw data and the LASSO method in Section 3.3, but show other relevant results in the figures below for comparison.

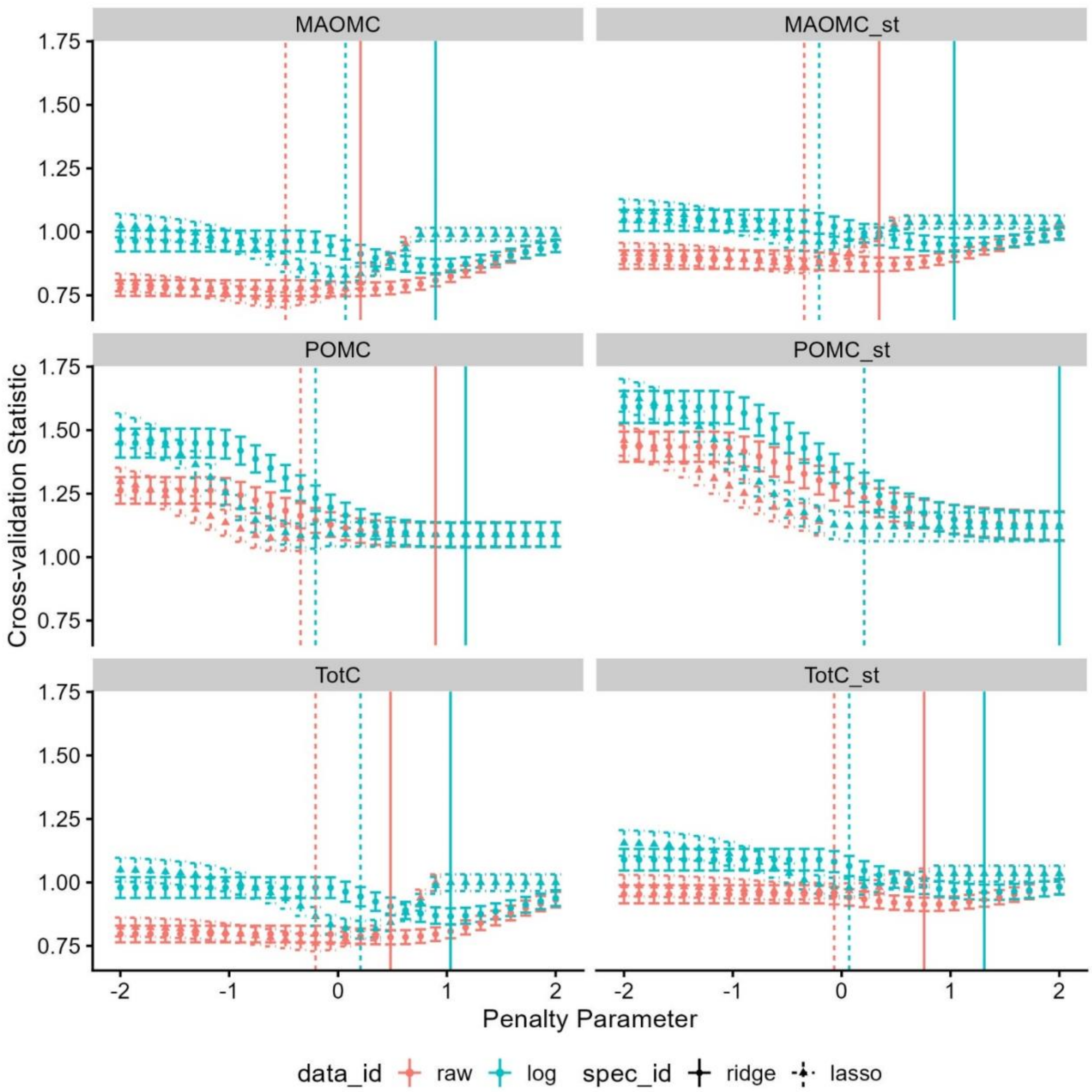


*Figure S8. Regularization curves used to select the penalty parameter for the regression results in Section 3.3. For each penalty parameter value, we performed 10-fold cross validation repeated 10 times (100 regressions per parameter) to get an estimate of the predictive performance. The vertical lines indicate the optimal penalty parameter based on root mean square error (rmse) to select the trained model. Note that the root mean square error is normalized by the standard deviation of the test set, so that a value greater than 1 indicates that the model is worse than the mean in a predictive sense. Here, we fit a model with and without log-transformed predictors as well both ridge and LASSO regularization methods for both fractionated and total SOC concentration (left column) and stocks (right column). These results are for the "root" predictors only.*

Once we tuned the penalty hyper-parameter, we next estimated the sampling distribution of the effects and predictive performance rather than simply fitting tuned model to full data set for a point estimate. To do this, we repeated the 10-fold cross-validation 20 times using the tuned penalty parameter, which produced 200 estimates of the out-of-sample performance (Fig. S9) and predictor effects (Fig. S10–11). Note this is very similar to bootstrapping, except we guarantee the model is trained on 90% of the data rather than the average 63% in bootstrapping. This is critical in our analysis as we only have 69 data points. As in the hyperparameter tuning, we observed that POM has very poor predictive performance (Fig. S9), which results in uncertain predictor effects centered around 0 (Fig. S10–11).

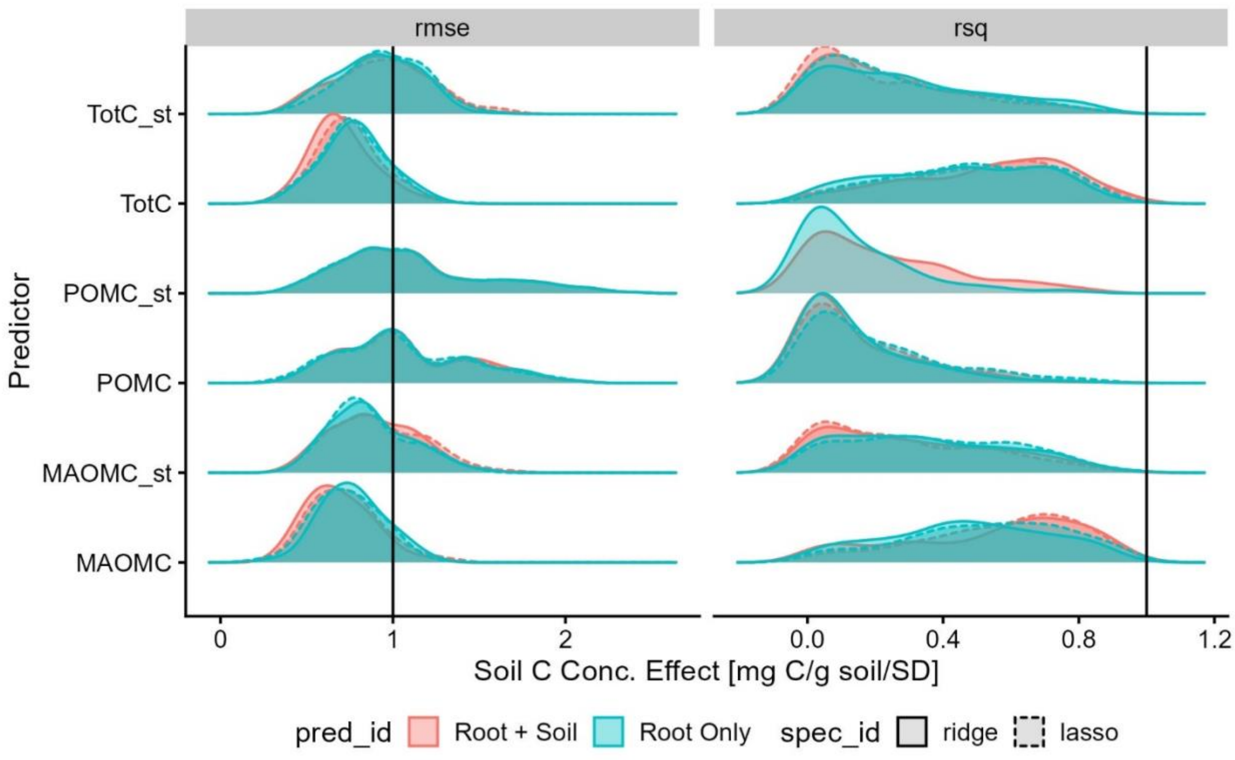


*Figure S9. The estimated sampling distribution of model performance for the tuned regression models in Fig. S8. Here, we performed 10-fold cross validation repeated 20 times (200 samples) and gathered the tuned model performance on the out-of-sample set (10% of the data). Note that the root mean square error is normalized by the standard deviation of the test set, such that a value greater than 1 indicates that the model is worse than the mean in a predictive sense.*

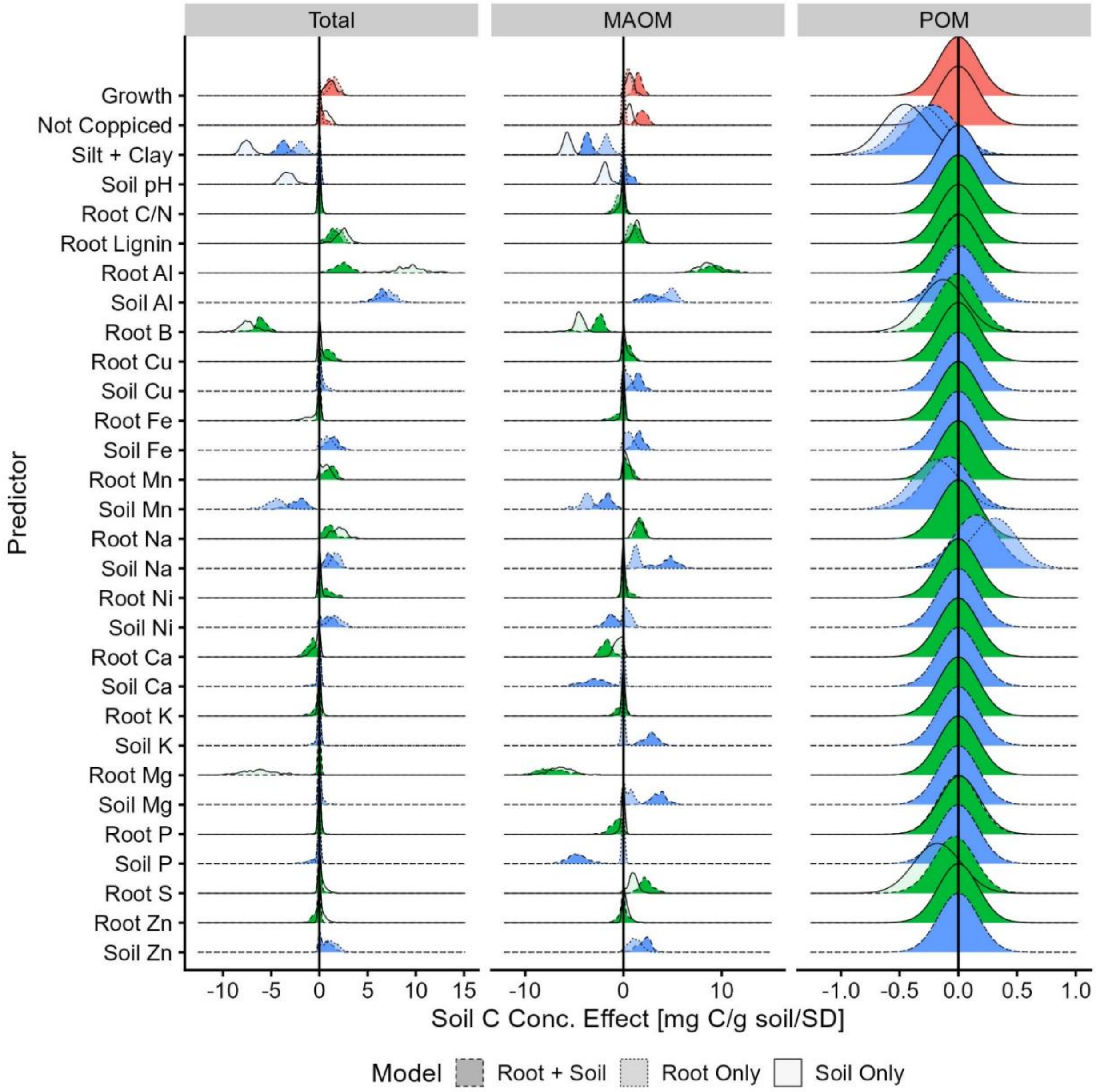


*Figure S10. The regularized regression effects on C concentration in various soil fractions for all three predictor sets tested. Total C and MAOM-C has similar effect sized. The regression model poorly fits the POM-C data (Fig. S9), as indicated in this figure by the large spread in effects centered around 0 for most phenotypes.*

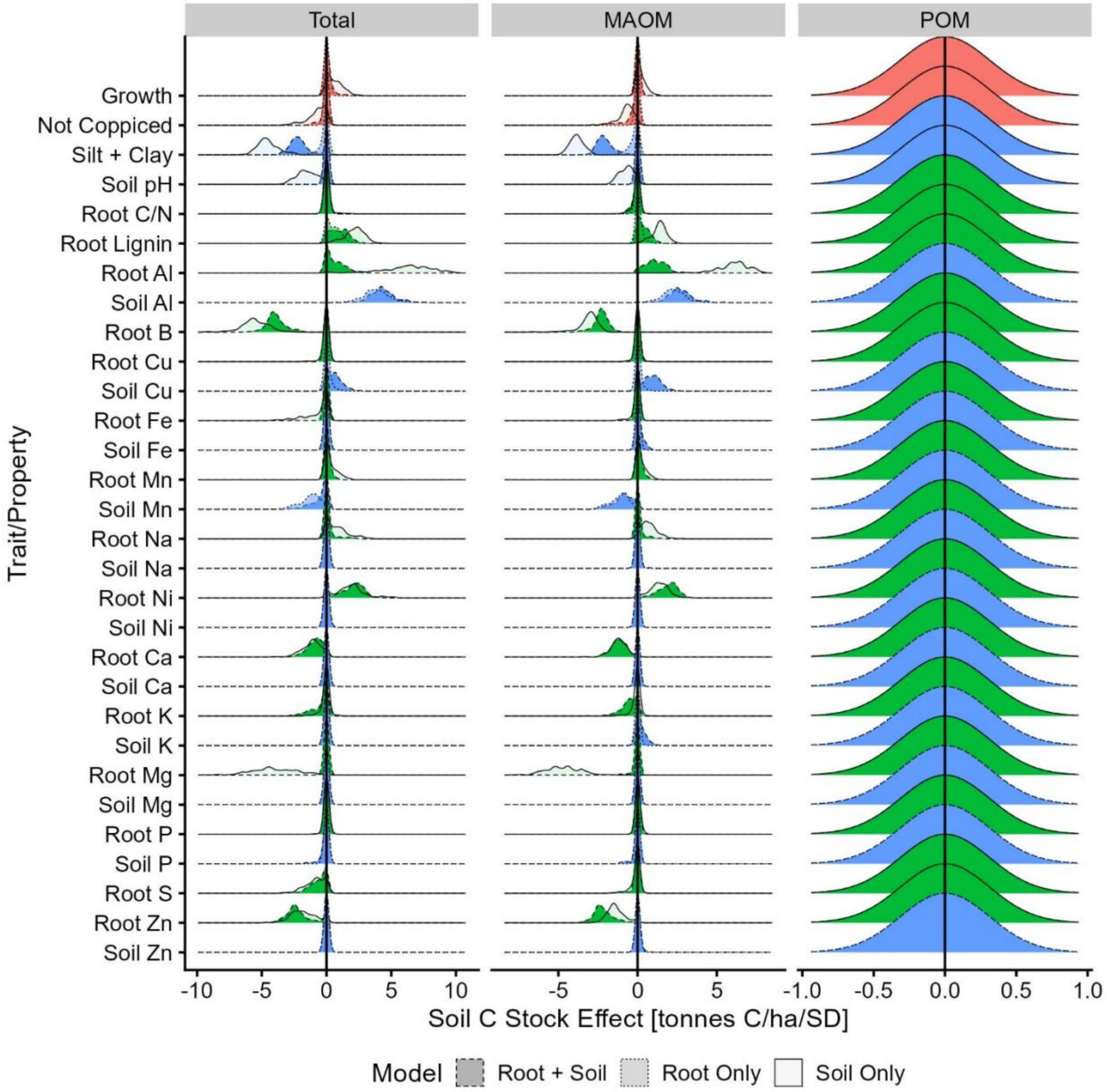


*Figure S11. Same as Fig. S10 except for soil C stocks rather than concentrations.*
